# ADA2 is a lysosomal DNase regulating the type-I interferon response

**DOI:** 10.1101/2020.06.21.162990

**Authors:** Ole Kristian Greiner-Tollersrud, Vincent Boehler, Eva Bartok, Máté Krausz, Aikaterini Polyzou, Johanna Schepp, Maximilian Seidl, Jan Ole Olsen, Cristian R. Smulski, Salvatore Raieli, Katrin Hübscher, Eirini Trompouki, Regina Link, Hilmar Ebersbach, Honnappa Srinivas, Martine Marchant, Dieter Staab, Danilo Guerini, Sebastian Baasch, Philipp Henneke, Georg Kochs, Gunther Hartmann, Roger Geiger, Bodo Grimbacher, Max Warncke, Michele Proietti

## Abstract

Deficiency of adenosine deaminase 2 (DADA2) is a severe, congenital syndrome, which manifests with hematologic, immunologic and inflammatory pathologies. DADA2 is caused by biallelic mutations in *ADA2*, but the function of ADA2, and the mechanistic link between ADA2 deficiency and the severe inflammatory phenotype remains unclear. Here, we show that monocyte-derived proteomes from DADA2 patients are highly enriched in interferon response proteins. Using immunohistochemistry and detailed glycan analysis we demonstrate that ADA2 is post-translationally modified for sorting to the lysosomes. At acidic, lysosomal pH, ADA2 acts as a novel DNase that degrades cGAS/Sting-activating ligands. Furthermore, we define a clear structure-function relationship for this acidic DNase activity. Deletion of ADA2 increased the production of cGAMP and type I interferons upon exposure to dsDNA, which was reverted by ADA2 overexpression or deletion of *STING.* Our results identify a new level of control in the nucleic acid sensing machinery and provide a mechanistic explanation for the pathophysiology of autoinflammation in DADA2.

**One Sentence Summary:** ADA2 is a lysosomal nuclease controlling nucleic acid sensing and type I interferon production.

## Introduction

Deficiency of adenosine deaminase 2 (DADA2) is an autosomal recessive disease characterized by a defective regulation of the inflammatory response (*1*, *2*). The disease was first described in 2014 in young patients with a polyarteritis nodosa-like phenotype and biallelic mutations in the *ADA2* gene (previously called *CECR1*). To date, more than 200 DADA2 patients have been reported, and the description of the clinical phenotype has expanded (*1*–*5*). DADA2 shares clinical and laboratory signs with type I interferonopathies, a group of genetically heterogeneous disorders characterized by autoinflammation and constitutive activation of the antiviral type I interferon (IFN) response. However, how ADA2 regulates the immune response is currently unknown (*6*).

Adenosine deaminases (ADA) are generally accepted to be essential for the metabolism and neutralization of purine nucleosides, respectively ADA1 within the intracellular and ADA2 within the extracellular space. However, in comparison to ADA1, ADA2 has a low affinity for adenosine (Ado) and 2’-deoxyadenosine (dAdo) (Km ~ 2 mM) and the adenosine concentration in the extracellular space is 10,000 fold less than its Km value (~ 0.2 μM) (*7*–*10*). Accordingly, the biological relevance of the ADA2 deaminase activity is a matter of current debate. Furthermore, while ADA1 is ubiquitously expressed (*11*), ADA2 is preferentially expressed in monocytes and macrophages (*12*–*14*) (Fig. 1A-B). Altogether these observations raise the possibility that ADA2 has a distinct function, unrelated to its adenosine deaminase activity.

**Fig. 1:**
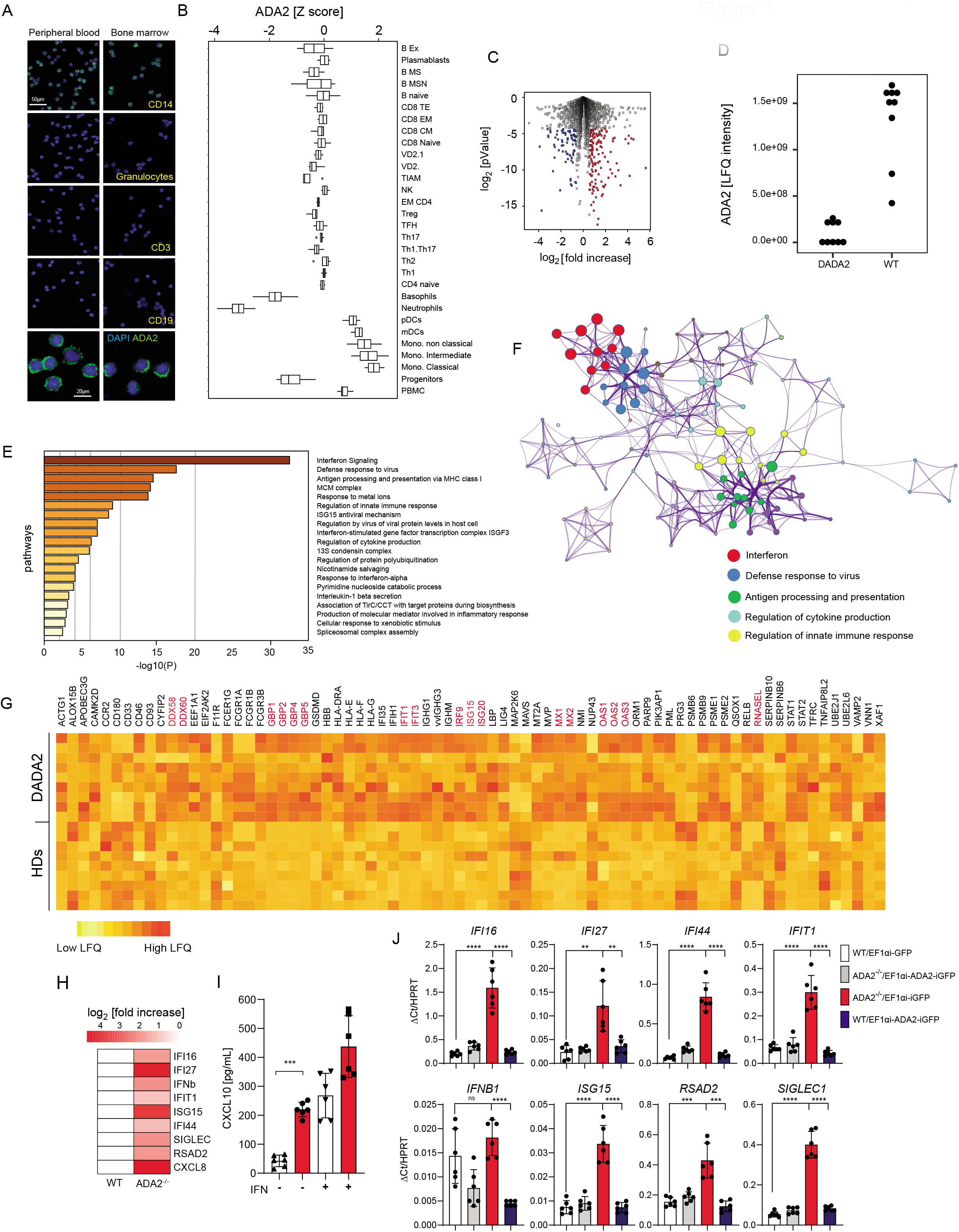
DADA2 proteomic signature of peripheral monocytes. (A) ADA2 expression in CD19^+^, CD3^+^, neutrophils and CD14^+^ immune cell subtypes FACS sorted from peripheral blood or bone marrow of healthy donors detected by immunofluorescence. (B) Analysis of ADA2 expression in different immune cells subtypes analyzed from GSE107011 RNAseq dataset (*34*). (C) Volcano plot showing differentially regulated (red=up-regulated, blue=down-regulated) proteins in monocytes proteomes of DADA2 patients in comparison to age and sex-matched healthy donors. (D) ADA2 protein amount, as measured by Label Free Quantification (LFQ intensity) in proteomes from DADA2 patients (left) and healthy controls (right). (E) Pathway analysis showing enrichment of interferon and related pathways in DADA2 proteomes compared to controls. (F) Interaction network on the pathways analysis above. (G) Heatmap showing proteins differentially expressed in DADA2 monocytes with respect to controls. In red are highlighted proteins involved in the interferon response (H) RNA expression of 9 interferon induced genes as measured by realtime PCR in *ADA2^-/-^* and WT THP-1 cells. Heatmap is showing the log2 of the fold change (*ADA2^-/-^* vs WT ratio). (I) Measurement of CXCL10 concentration in the supernatant of *ADA2^-/-^* and WT THP-1 cells measured by ELISA. The lines over the bars show the statistical comparisons reported in the figure (3 independent experiments, each experiment with duplicates; test used, unpaired two tailed t test with Welch’s correction, *** = p <= 0.001). (J) RNA expression of 8 interferon induced genes as measured by realtime PCR in *ADA2*^-/-^ and WT THP-1 cells transduced with lentiviruses carrying human ADA2 or the corresponding empty lentivirus. Y axis shows the log. of the Delta Ct relative to HPRT. The lines over the bars show the statistical comparisons reported in the figure (3 independent experiments, each experiment with duplicates; test used, unpaired two tailed t test with Welch’s correction, ** = p <= 0.001, *** = p <= 0.001, **** = p <= 0.0001).

To gain insight into the molecular consequences of ADA2 deficiency, we compared the proteomes of purified monocytes from six patients with different biallelic mutations in *ADA2* (suppl. Fig. 1A-B) to sex- and age-matched controls. At a false discovery rate of 1%, we identified more than 10,000 different proteins with an average of 7,500 proteins per sample. 138 proteins were significantly (log2 fold change cut-off of 1.5 and Welch’s test p-value <=0.05) upregulated and 79 proteins were downregulated in DADA2 patients compared to healthy individuals (Fig. 1C and Table S1). As expected, ADA2 was undetectable or strongly reduced in all proteomes from DADA2 patients (Fig. 1D). Pathways analysis of differentially expressed proteins revealed a remarkable up-regulation of interferon signaling-related pathways in DADA2 patients compared to controls (Fig 1E). Network analysis suggested that ADA2 deficiency mostly affected the innate immune response to viral infection (Fig. 1F). Differentially-expressed proteins comprised several interferon-stimulated genes (ISGs) (18,9), including guanylate binding proteins (GBP5, GBP4, GBP1 and GBP2), interferon-induced proteins with tetratricopeptide repeats (IFIT1 and IFIT3), 2’-5’-oligoadenylate synthase 3 (OAS3), signal transducer and activator of transcription 1 alpha/beta (STAT1), and the interferon-induced GTP-binding protein Mx1 (Fig. 1G). Proteins previously associated with interferonopathy (8–9, 19–21), were expressed at regular levels (Fig. S1C), supporting the growing consensus that DADA2 is a distinct clinical entity (*15*). Comparing ADA2null (no ADA2 protein detectable) and ADA2dim (residual ADA2 protein detectable) patients, we observed a significant (unpaired t.test < 0.01 & nonlinear fit R Square > 0.9) correlation between the ADA2 expression and the interferon signature (Fig. S2A). These data suggested that ADA2 directly controls the antiviral state acting as a rheostat of the type I interferon signature in monocytes.

To address whether the observed interferon signature was cell intrinsic, we generated an ADA2-deficient monocytic cell line (THP-1) (Fig. S3A-b). Wild-type and ADA2-deficient cells were subsequently tested by real-time-PCR for the expression of several interferon induced genes (ISGs), namely *IFI16, IFI27, IFI44, IFIT1, IFNB1, ISG15, RSAD2, SIGLEC1* and *CXCL8* (*16*). As shown in Fig. 1H, ADA2-deficient cells showed increased expression of all the measured ISGs. We also observed increased levels of secreted CXCL10, another interferon-induced protein, in the supernatants of ADA2-deficient THP-1 (Fig. 1I). Importantly, ADA2 reexpression by lentiviral transduction completely reverted the phenotype (Fig. 1L). Collectively, these data indicate that ADA2 negatively regulates the type 1 interferon pathway in monocytes in a cell-intrinsic manner.

Current consensus considers ADA2 to be a predominantly extracellular protein (*17*, *18*). However, our monocyte data above suggested an intrinsic function for ADA2, inconsistent with an extracellular adenosine deamination. We were particularly interested in understanding the cellular and subcellular expression pattern on ADA2 within secondary lymphoid organs, in which we (*19*) (Fig. S4A-F) have observed significant histopathological abnormalities in DADA2 patients. To address this question, we stained human tonsils for ADA2. ADA2 signal was confined to germinal centers and was mostly expressed in tingible body macrophages (Fig. S4 G, H). Interestingly, at high magnification, ADA2 was observed in vesicular structures resembling phagolysosomes (Fig. 2A). To address whether ADA2 was indeed present in lysosomal vesicles, we additionally stained tonsil sections with LAMP-1, a marker for lysosomal membranes. Immunofluorescence confirmed that ADA2 is contained within the lumen of LAMP1-positive lysosomal vesicles (Fig. 2B and inlet).

**Fig. 2:**
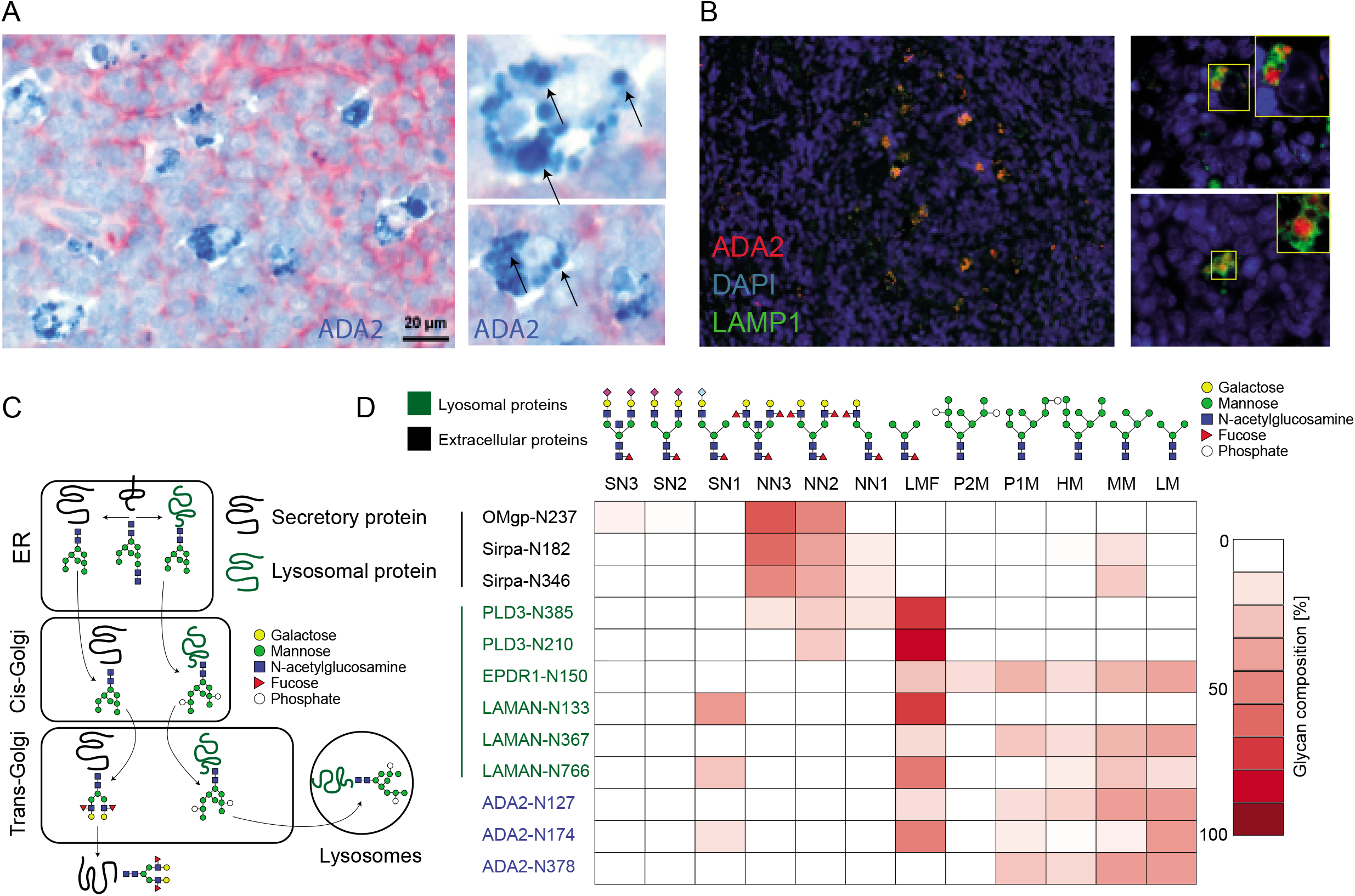
Localization of ADA2 in the lysosomes of tingible body macrophages and ADA2 lysosomal targeting via the mannose-6-phosphate dependent pathway. (A) ADA2 staining (Blue) in tingible body macrophages. On the right panels the arrows indicate ADA2 positive cellular organelles resembling phagolysosome (B) Immunofluorescence of human tonsils showing LAMP1 positive lysosomes containing ADA2, the inlets show higher (40X) magnification; (C) Diagram showing lysosomal sorting of Mannose-6-phosphorylated proteins (D) Heatmap showing the comparison of the glycan structures linked to specific glycosylation sites of pbADA2 and to other known lysosomal (green) and extracellular (black) proteins. All the proteins have been purified from porcine brain. The color scale represents the percentage of the identified glycan structures (Fig. S4) linked to the indicated amino acid residues.

In order to be targeted to the lysosome, soluble proteins typically need to be tagged with mannose-6-phosphate (M6P) on a subset of their N-glycans, which are recognized by the mannose-6-phosphate sorting receptor in the trans-Golgi (*20*) (Fig. 2C). Once in the lysosomes, these glycan structures are intensively trimmed by lysosomal phosphatase and exoglycosidases. To confirm that ADA2 is a lysosomal protein and to understand how it is targeted to these cellular organelles *in vivo*, we analyzed the glycosylation pattern of endogenous ADA2. Except in rodents, ADA2 orthologs are present in all mammals, including pigs, which exhibits a high similarity with human ADA2 (Fig. S5A-B). Further, analysis of public datasets showed that ADA2 is highly expressed by microglia (*21*) (Fig. S5C). Thus, to access sufficient material for glycoanalysis, we purified ADA2 from porcine brain (Fig. S6A-D). Data are available via ProteomeXchange (*22*) with identifier PXD019373. We found that endogenous ADA2 contains a high proportion of M-6 phosphorylated glycans linked to three of its glycosylation sites (N127, and N378). The spectrum of glycan structures in ADA2 overlaps with the lysosomally-trimmed glycans of known lysosomal proteins, such as lysosomal alpha-mannosidase (LAMAN), phospholipase D3 (PLD3) and ependymin related protein 1 (EPDR1), and it is different from the glycan structures of the extracellular proteins oligodendrocyte myelin glycoprotein (OMgp) and signal regulatory protein alpha (Sirpa) (Fig. 2D). Altogether, these data indicate that in monocytes/macrophages ADA2 is targeted to the lysosomes *via* the M6P-dependent pathway.

Since the adenosine deaminase activity of ADA2 is incompatible with the acidic-lysosomal pH (*18*), these results reinforced the notion that in the lysosome ADA2 serves a distinct function, unrelated to its deaminase activity. In agreement with this, there are clear differences in the clinical phenotypes of ADA1 *vs*. ADA2 deficiency (Table S2), while DADA2 shares clinical and immunological similarities with other type I interferonopathies (*23*–*25*). Of particular relevance to this study, there is a substantial clinical overlap of DADA2 with DNase type II deficiency (*24*) (Table S2), raising the question of whether ADA2 is involved in DNA sensing.

To search for a new function of the protein we evaluated the differences between the active site of ADA1 and ADA2 (*16*). Although both enzymes accommodate a purine in the active site pocket, ADA2 possesses an enlarged pocket allowing the positioning of a purine-linked ribose unit (*17*, *18*). This “widening” of the active site may allow a larger nucleotide-containing molecule, such as DNA or RNA, to serve as a substrate for ADA2. This observation, together with the clinical similarities between DADA2 and DNase2 deficiency, led us to hypothesize that ADA2 could possess a hitherto undescribed DNase function. Furthermore, given its lysosomal localization, we considered the possibility that a putative DNase activity of ADA2 could be pH dependent. Accordingly, when we incubated endogenous porcine ADA2 with plasmid DNA, the plasmid was degraded with a pH optimum between 4.6 and 5, corresponding to the lysosomal pH (Fig. 3A), but not at neutral pH. More detailed investigations revealed that, at low concentrations of ADA2, the supercoiled plasmid DNA was converted into a relaxed circular molecule, while at a higher concentration of ADA the relaxed circular plasmid was converted into a linear form, and smaller DNA fragments appeared (Fig 3B). These results suggest that ADA2 acts as a nicking endonuclease. Any contamination with the only currently known lysosomal DNase, DNase 2, was ruled out by the separation of ADA2 from DNase 2 on hydroxyapatite chromatography (Fig. S4B). Data are available via ProteomeXchange (*22*) with identifier PXD019373.

**Fig. 3:**
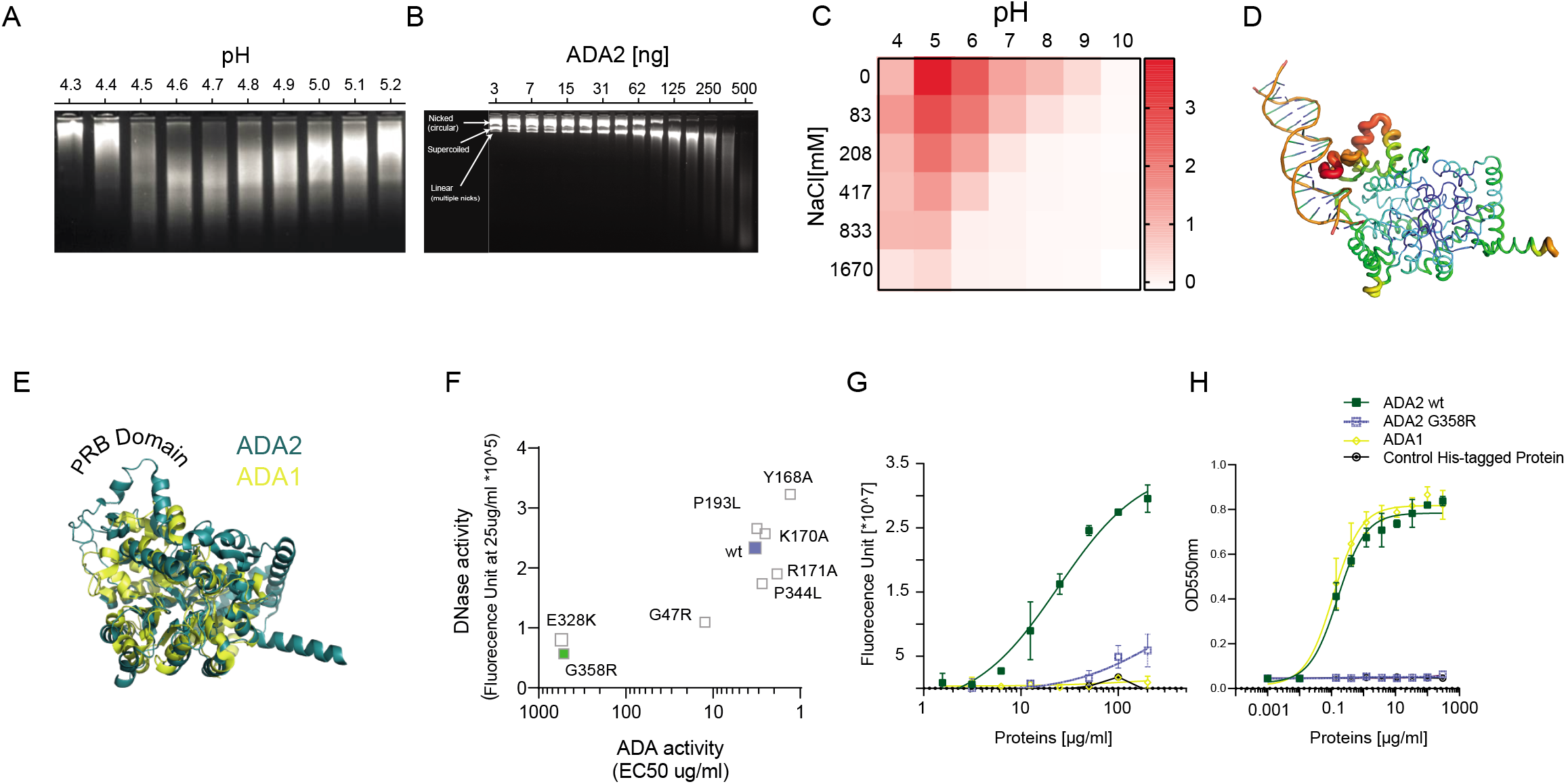
Acidic DNase activity of porcine and human ADA2. (A-B) Gel electrophoresis showing DNA degradation after 20’ incubation of plasmid DNA with pbADA2 at different pH (A) or at pH4.5 and different protein concentrations (B). (C) DNase activity of recombinant human ADA2 (25μg/ml) as measured by FRET assay evaluated at 20’ and at different pH and NaCl concentration. (D) Modeling of the binding between dsDNA and human ADA2 (PDB entry: 3LGD) (*17*). The top scored model corresponds to the interaction of the dsDNA molecule with the PRB domain, which is located close to the active site and therefore places the DNA molecule on top of the catalytic cleft. (E) Comparison between the ADA1 (PDB entries: 3IAR) (*35*), and ADA2 (3LGD) structure showing the absence of a ~40 amino acids insertion (aa 111-147), corresponding to the PRB domain in ADA1 in comparison to ADA2. (F) Evaluation of DNase and ADA activities of 8 recombinant ADA2 mutants and correlation. In green is reported the mutant (G358R) selected for further large-scale validation, in blue the human wild type ADA2. (G-H) Evaluation of DNase and ADA activities of the recombinant G358R mutant, ADA1 and ADA2 and the control His-tagged protein.

To validate these observations, we expressed and purified recombinant human ADA2 from HEK cells and measured its nuclease activity using a FRET-based assay. This assay confirmed the DNase activity of recombinant ADA2, as well its maximal activity at lysosomal pH (Fig. 3C). The DNase activity of ADA2 inversely correlated with the salt concentration (Fig. 3C), similar to DNase type I (*26*). Importantly, MS/MS analysis of our recombinant ADA2 did not detect any contamination by known nucleases (Fig. S7A). Data are available via ProteomeXchange (*22*) with identifier PXD019382. Modeling of the binding of ADA2 with double-stranded DNA suggested that the a ~ 40 amino acid domain called PRB domain (Fig. 3D), which is present in ADA2, but absent in ADA1 (*26*), mediates the interaction between the protein and the dsDNA molecule (Fig. 3E). In agreement, ADA1 lacked nuclease activity (Fig. 3G, H). These results provide a mechanistic explanation of the stark clinical differences between ADA1 and ADA2 deficiency.

To gain further insight into the structure-function relationship between ADA2 and DNase activity, we selected variants of ADA2 that cause defined clinical phenotypes (*27*) and were predicted to be important for ADA2 DNase activity by structural modeling (Table S.3 and Fig. S7B). These variants were then expressed in HEK cells, purified, and tested for DNase and deaminase activity. Of note, many of the single point mutants and domain deletions, including the PRB domain, showed a severe loss of expression, indicating that these residues are essential for the protein stability (Table S.3). Nevertheless, eight mutants had a sufficient expression for further analysis of enzymatic activity. Evaluation of adenosine deamination *versus* DNase activity revealed a strong correlation between the two (Fig. 3F), with the adenosine deamination correlating well with values reported previously for these variants (*27*) (Fig. 3F). To further validate these findings, we selected ten variants for larger scale production and stringent purification. The stringently purified proteins fully confirmed the results from the smaller scale screen (Fig. S7C). Importantly, we detected variants, like G358R, which showed absent or largely reduced ADA (*27*) and DNase activity. (Fig 3G-H). The G358R amino acid change replaces an highly conserved glycine with arginine at position 358 of the ADA2 protein (p.Gly358Arg) and it has been reported in homozygosity or in combination with another ADA2 variant (compound heterozygosity) in individuals with DADA2 and bone marrow pathology (*27–30*).

Collectively, our data suggest that ADA2 deficiency could lead to a defective degradation of double-stranded DNA, which would be expected to provide more ligands for DNA-sensing machinery, in particular the cGAS-STING pathway. Alternatively, ADA2 deficiency may more generally elicit monocytes sensitivity to type I interferon stimulants, *via* an unknown mechanism unrelated to its DNase activity. To directly address these hypotheses, we measured type I IFN in the supernatants of ADA2 KO monocytic cells following stimulation with different pattern recognition receptor agonists. Specifically, cells were stimulated with: Poly-IC, LPS, R-837 or interferon stimulatory DNA. Poly-IC is a synthetic RNA analogous sensed by three receptors: TLR3, that signals through adapter Toll/IL-1R domain–containing adapter-inducing IFN-β (TRIF); and two cytoplasmic receptors, retinoic acid–inducible gene I (RIG-I) and melanoma differentiation–associated protein-5 (MDA-5), which both signal through mitochondrial antiviral-signaling protein (MAVS); LPS, is an agonist of the TLR4, which signals through MyD88; R-837 is, an imidazoquinoline amine analog to guanosine, sensed by the TLR7; interferon stimulatory DNA, a dsDNA oligomer that, when transfected into various cell types, including macrophages, activates the cGAS-STING pathway (*31*). ADA2 deficiency did not affect type I interferon production in response to Poly-IC, LPS or R-837 (Fig. 4A). In contrast, we observed an increased induction of interferon when ADA2-deficient cells were exposed to interferon stimulatory DNA (Fig. 4B). This difference was reverted by the over-expression of wild-type ADA2 and by the genetic deletion of STING (Fig. 4B). These data strongly suggest that ADA2 deficiency determines a defective sensing of dsDNA and a subsequent hyperactivation of STING signaling. In agreement with this, STING deletion also completely reverted the interferon signature (Fig. 4C) and CXCL10 secretion (Fig. 4D) in ADA2 KO monocytic cells. Deletion of the critical TLR adaptor protein MyD88 or the intracellular dsRNA sensor MAVS did not affect the interferon signature of ADA2-deficient cells, further supporting the hypothesis that this interferon response is dsDNA-cGAS-STING-dependent (Fig. 4C, D). STING is activated by cGAMP, a cyclic dinucleotide that is produced by cGAS upon DNA stimulation (*32*). Thus, intracellular cGAMP levels correlate with the level of dsDNA detected by cGAS (*33*). We therefore measured intracellular cGAMP in wild-type and ADA2 KO monocytic cells following stimulation with dsDNA. As shown in Fig. 4E, ADA2 deficiency was associated with a significant increase in the production of cGAMP, which could be reverted by the over-expression of ADA2 (Fig. 4E).

**Fig. 4:**
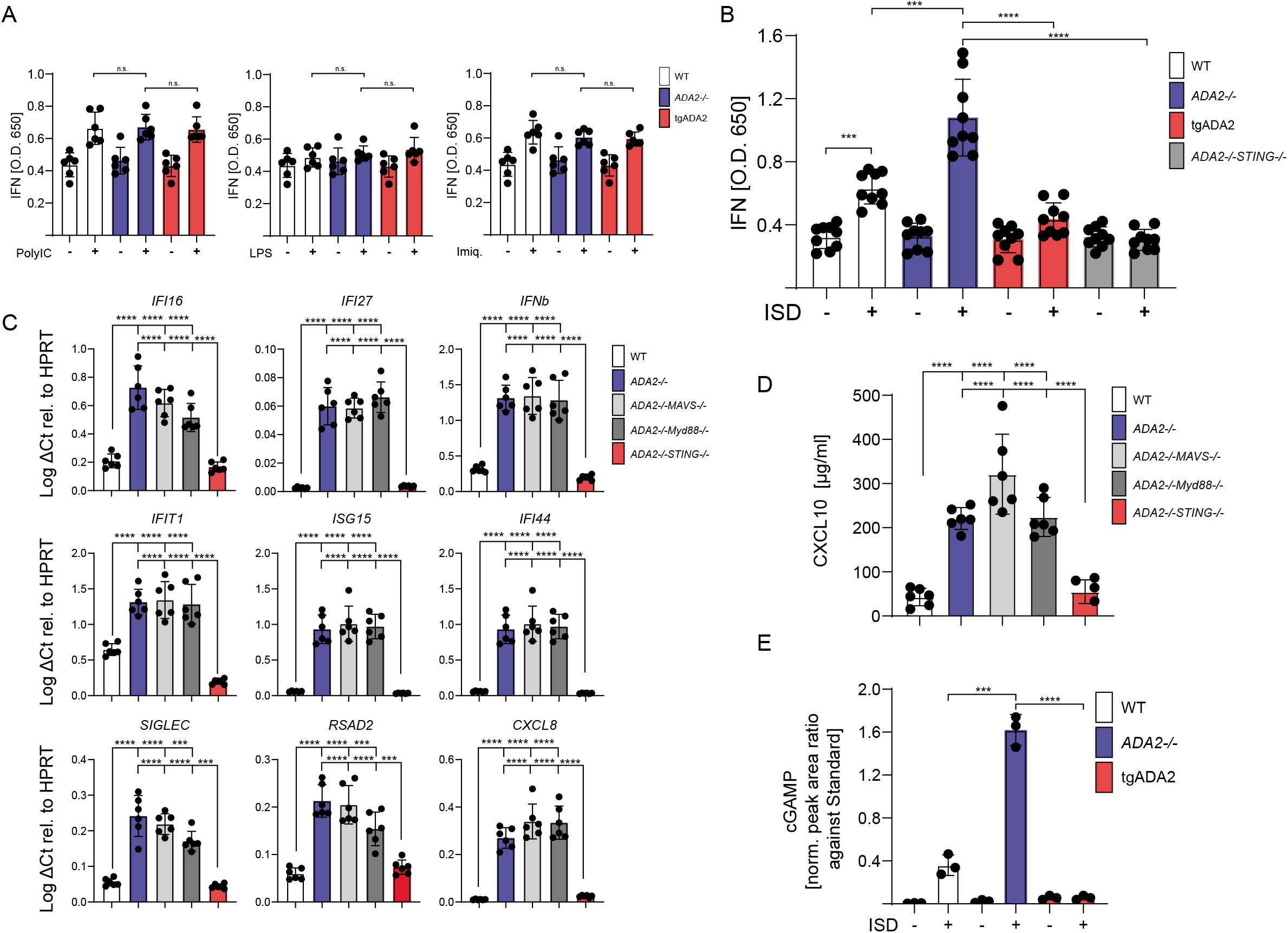
ADA2 dependent cGAS/STING signaling. (A) Concentration of type I interferon, as measured by HEK-Blue IFN-α/β report assay, in the supernatant of WT (white), *ADA2^-/-^* (blue) and *ADA2^-/-^* lentiviral transduced with human ADA2 (red) THP-1 cells, after overnight stimulation with PolyIC, LPS or Imiquimod. The lines over the bars show the groups compared for statistical analysis (2 independent experiments, each experiment with triplicates; test used, unpaired two tailed t test, n.s = not significant). (B) Concentration of type I interferon, as measured by HEK-Blue IFN-α/β report assay, in the supernatant of WT (white), *ADA2^-/-^* (blue) and *ADA2^-/-^* lentiviral transduced with human ADA2 (red) and *ADA2*^-/-^ *STING*^-/-^ (gray) THP-1 cells, after overnight stimulation with Interferon Stimulating DNA (ISD). The lines over the bars show the statistical comparisons reported in the figure (3 independent experiments, each experiment with triplicates; test used, unpaired two tailed t test with Welch’s correction, *** = p <= 0.001, **** = p <= 0.0001). (C) Expression of 9 interferon stimulated genes measured by RT-PCR in unstimulated WT, *ADA2*^-/-^, *ADA2*^-/-^*STING*^-/-^, *ADA2^-/-^MAVS^-/-^, ADA2^-/-^Myd88^-/-^* THP-1 cells. Y Axis shows the log of the Delta Ct relative to HPRT. The lines over the bars show the groups compared for statistical analysis (3 independent experiments, each experiment with duplicates; test used, unpaired two tailed t test with Welch’s correction, *** = p <= 0.001, **** = p <= 0.0001). (D) Concentration of CXCL10 in the supernatant of unstimulated WT, *ADA2^-/-^, ADA2^-/-^STING^-/-^, ADA2^-/-^MAVS^-/-^, ADA2^-/-^Myd88^-/-^* THP-1 cells, measured by ELISA (3 independent experiments, each experiment with duplicates; test used, unpaired t test with Welch’s correction, *** = p < 0.001, **** = p < 0.0001). (E) cGAMP production in WT, *ADA2*^-/-^ and *ADA2*^-/-^ lentiviral transduced with human ADA2 after exposure to dsDNA.

In summary, our discovery of the lysosomal localization and the unique structure of ADA2 together with the severe interferonopathy observed in DADA2 patients with clinical similarities to DNase2 deficiency led us to the discovery of a hitherto unappreciated DNase function of ADA2. This function has presumably escaped prior detection due to its unique dependence upon acidic, lysosomal pH. Building on the DNase function, we have established that ADA2 is essential for the tuning of nucleic acid sensing in myeloid cells with a remarkable specificity for the cGAS-STING pathway. These data strongly suggest that altered degradation of dsDNA is the primary driver of the interferonopathy observed in DADA2 patients. Our results reveal a novel level of control in nucleic acid sensing and not only provide important mechanistic insight into the pathogenesis of ADA2 deficiency and putatively more common and complex autoinflammatory conditions such as lupus erythematosus or vasculitis, but additionally opens the horizon for the development of personalized treatment approaches for these conditions.

## Acknowledgments

We are deeply grateful to all affected individuals, their families and all healthy donor controls who participated in this study. We thank Pavla Mrovecova, Mary Buchta, Hanna Haberstroh, Nadine Glaser (CCI, Uniklinik Freiburg), and Thi-Thanh-Thao Tran (Novartis) for their excellent technical assistance. We thank Prof. Dr. Martin Schwemmle for providing SC35M encoding GFP. We thank Prof. Dr. Stephan Ehl (CCI, Uniklinik Freiburg), Ben Roediger (Novartis) and Dr. Elisabetta Traggiai (Novartis) for helpful advice and discussion. Some samples have been taken from the CCI-biobank, a partner of the Freeze Biobank Freiburg.

## Author contributions

Conceptualization: O.K.G.T., B.G., M.W., E.B., C.R.S, G.H, M.H., and M.P.. Methodology: O.K.G.T., R.G., C.R.S., M.S., A.P. M.W. and M.P.. Validation: O.K.G.T., M.W., B.G. and M.P.. Formal Analysis: M.P., M.W. and A.P.. Investigation: O.K.G.T., M.K., M.S., S.B., C.R.S., M.S., K.H., J.S., H.E., S.B., M.M., V.B., M.B., D.S., N.J.G, D.G., M.W. and M.P. Resources: O.K.G.T., M.W., B.G., J.S., A.P. and M.P.. Data Curation: A.P, M.W. and M.P. Writing: O.K.G.T., M.W., B.G., and M.P.. Visualization: O.K.G.T., M.W. and M.P.. Supervision: M.P. O.K.G.T., B.G., M.W.. Funding Acquisition: O.K.G.T., E.T., B.G. and M.P.

## Supplementary Materials

### Materials and Methods

#### Monocytes isolation from peripheral blood mononuclear cells

Peripheral Blood Mononuclear Cells (PBMC) were prepared by centrifugation on a Ficoll gradient (Lymphoprep ™). Blood CD14^+^ monocytes were isolated from DADA2 patients or healthy donors’ PBMC by positive selection using magnetic beads (Miltenyi). Monocytes purity was 95%–98% as measured by flow cytometry (see suppl. Fig. 1A).

#### Sample Preparation for Proteome MS Analysis

Samples were processed as described by Rieckmann JC et al (*1*). In brief, cell pellets were washed with PBS and lysed in 8M urea, 10 mM HEPES (pH 8), 10 mM DTT. Cell pellets were sonicated at 4°C for 15 min (level 5, Bioruptor, Diagenode). Alkylation was performed in the dark for 30 min by adding 55 mM iodoacetamide (IAA). A two-step proteolytic digestion was performed. First, samples were digested at room temperature (RT) with LysC (1:50, w/w) for 3h. Then, they were diluted 1:5 with 50 mM ammoniumbicarbonate (pH 8) and digested with trypsin (1:50, w/w) at RT overnight. The resulting peptide mixtures were acidified and loaded on C18 StageTips. Peptides were eluted with 80% acetonitrile (ACN), dried using a SpeedVac centrifuge (Savant, Concentrator plus, SC 110 A), and resuspended in 2% ACN, 0.1% trifluoroacetic acid (TFA), and 0.5% acetic acid.

#### LC-MS/MS for Analysis of Proteomes

Peptides were separated on an EASY-nLC 1200 HPLC system (Thermo Fisher Scientific) coupled online to a Q Exactive mass HF spectrometer via a nanoelectrospray source (Thermo Fisher Scientific). Peptides were loaded in buffer A (0.1% formic acid) on in house packed columns (75 μm inner diameter, 50 cm length, and 1.9 μm C18 particles from Dr. Maisch GmbH). Peptides were eluted with a non-linear 180 min gradient of 5%–60% buffer B (80% ACN, 0.1% formic acid) at a flow rate of 250 nl/min and a column temperature of 50°C. The Q Exactive was operated in a data dependent mode with a survey scan range of 300-1650 m/z and a resolution of 60’000 at m/z 200. Up to 10 most abundant isotope patterns with a charge ≥ 2 were isolated with a 1.8 Th wide isolation window and subjected to higher-energy C-trap dissociation (HCD) fragmentation at a normalized collision energy of 27. Fragmentation spectra were acquired with a resolution of 15,000 at m/z 200. Dynamic exclusion of sequenced peptides was set to 30 s to reduce the number of repeated sequences. Thresholds for the ion injection time and ion target values were set to 20 ms and 3E6 for the survey scans and 55 ms and 1E5 for the MS/MS scans, respectively. Data were acquired using the Xcalibur software (Thermo Scientific).

#### Analysis of Proteomics Data

MaxQuant software (version 1.5.3.54) was used to analyze MS raw files (*2*). MS/MS spectra were searched against the human Uniprot FASTA database (Version Oct 2017, 71’775 entries) and a common contaminants database (247 entries) by the Andromeda search engine. Cysteine carbamidomethylation was set as fixed and N-terminal acetylation and methionine oxidation as variable modification. Enzyme specificity was set to trypsin with a maximum of 2 missed cleavages and a minimum peptide length of 7 amino acids. A false discovery rate (FDR) of 1% was required for peptides and proteins. Peptide identification was performed with an allowed precursor mass deviation of up to 4.5 ppm and an allowed fragment mass deviation of 20 ppm. Nonlinear retention time alignment of all measured samples was performed in MaxQuant. Peptide identifications were matched across different replicates within a time window of 1 min of the aligned retention times. A minimum ratio count of 1 was required for valid quantification events via MaxQuant’s Label Free Quantification algorithm (MaxLFQ). Data were filtered for common contaminants and peptides only identified by side modification were excluded from further analysis.

#### Bioinformatic analysis of proteomic data

Proteins with fold change cut-off 1.5 and p-value (Welch’s approach) <=0.05 were considered to be significantly differentially enriched. R/Bioconductor (version 1.0.1.) (*3*, *4*) and DEP package (*5*) were used for the differential enrichment analysis of the proteomics data. The volcano plot that was constructed to depict the significant deregulated proteins was generated with ggplot2 package (*6*). Gene ontology and pathway analysis were performed with DAVID knowledgebase(*7*), only pathways and biological processes with p-value <=0.05 were considered significantly enriched. Cytoscape (*8*) and Metascape (*9*) were used for network construction.

#### FACS sorting from PBMCs or bone marrow bone marrow cellular suspension

For sorting, either PBMCs or bone marrow cellular suspension were used. The latter were prepared by red blood cell lysis of bone marrow aspirates and subsequent washing with PBS. The cells were stained with the antibodies listed in supplementary table 1 and 2. At the sorting facility, the cells were subjected to a MoFlo Astrios EQ cell sorter. Post-sort controls on the same machine showed a purity of > 98 % for the sorted populations. B cells: lymphocytes FSC/SSC -> CD19^+^CD3^-^CD14^-^; Monocytes: monocytes FSC/SSC -> CD19^-^CD3^-^CD14^+^; T cells lymphocytes FSC/SSC -> CD19^-^CD3^+^CD14^-^; Granulocytes: granulocytes FSC/SSC.

Antibodies and relative working dilution used for FACS sorting of peripheral lymphocyte subpopulations

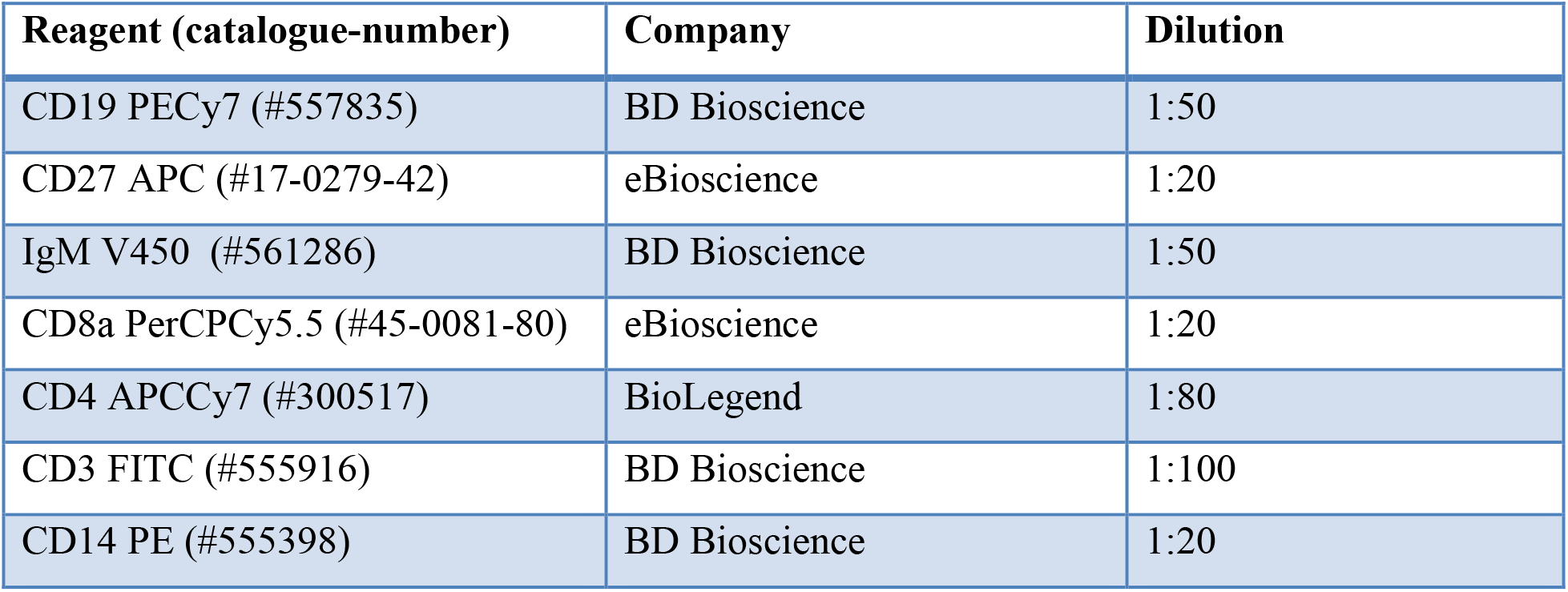

#### Microscopy

For histopathological sample analysis, 2μm sections were taken from buffered formalin fixed paraffin embedded biopsies and put on coated slides (SuperfrostPlus, Langenbrinck, Emmendingen, Germany). Staining was done with a primary antibody against CD14 (rabbit monoclonal antibody, clone EPR3653, BioGenex, Fremont, USA), no antigen retrieval was necessary. Secondary antibody was taken from the K5005 kit (Dako) according to the manufacturers guidelines and visualized with an alkaline phosphatase based red chromogen reaction from the K5005 kit (Dako). After denaturation of this first antibodies as described (Poc et al, manuscript in preparation), staining against ADA2 (anti CECR2 rabbbit polyclonal antibody, Sigma-Aldrich, Taufkirchen, Germany) was done after heat mediated epitope retrieval in a steamer for 30 minutes in TRIS buffered saline at pH6.1. The secondary antibody was again taken from the K5005 kit (Dako), visualized by an alkaline phosphatase based blue chromogen reaction (StayBlue/AP, Abcam, Cambridge, UK). Photos were taken with an Olympus BX 51 microscope (Olympus Hamburg, Germany) with the AxioCam MRc microscope camera (Carl Zeiss, Oberkochen, Germany). For immunofluorescence, the protocol by Poc et al was used again. The first staining step was done against LAMP1 (clone eBioH4A3, ThermoFisher, Darmstadt, Germany) after heat mediated epitope retrieval in a steamer for 30 minutes in TRIS buffered saline at pH6.1, visualized in the 488 channel (Alexa488, Thermofisher). The second staining step was done against ADA2 with the antibody as mentioned previously and high pressure cooking epitope retrieval for two minutes in citric buffer pH6, visualized in the 555 channel (Alexa555, Thermofisher). Nuclei were couterstained with DAPI 1:1000 (ThermoFisher). Images were taken with a fluorescence microscope (Axioplan 2, Zeiss) with the AxioCam MRm microscope camera. For ADA2 detection by Immunofluorescence, slides were prepared prior to cytospin as follows: They were soaked in NaOH-Ethanol solution and incubated with gentle rotation for 2 hours at room temperature in order to roughen the surface of the glass for better adhesion. Afterwards, they were washed extensively under running ddH2O for 30 minutes to remove all traces of NaOH. Then, they were coated with 10 μg/mL Poly-D-Lysin in PBS for 45 minutes. After careful removal of the coating solution, slides were let to dry and stored at room temperature until further usage. Purified granulocytes, monocytes, T and B cells from healthy donor EDTA blood or from healthy donor bone marrow aspirates were cytospinned at 700g for 5 minutes at room temperature with a Hettich Rotofiy 32A cytospin centrifuge. Cells were fixed with precooled 1:1 acetone/methanol fixative 20 minutes at −20°C. Afterwards the fixative was aspirated, and the sample washed three times in PBS for 5 minutes. Then, the cells were blocked with blocking solution (5% human serum in PBS) for 1.5 hours at room temperature. Hereafter, cells were immunostained with a rabbit anti-human-CECR1 (HPA007888 Sigma; 1:100) in blocking solution over night at 4°C; or with the isotype control at the same concentration (Purified Rabbit IgG, P120-101 Bethyl; 1:2500), respectively. The next morning, they were washed with PBS twice for 5 minutes and stained again simultaneously with an anti-rabbit IgG conjugated with Anti-rabbit IgG Alexa Fluor^®^ 555 (1:500) and Hoechst (1:1000 = 10 μg/ mL) in blocking solution for 30 minutes at room temperature in the dark. After two washes with PBS, Prolong^®^ Gold Anti-fade Reagent and coverslip slides were mounted on the slides. Then, images were aquired with a Zeiss LSM 710 and later analyzed with Zen imaging-software (Zeiss).

#### Modeling of dsDNA binding to ADA2 and comparison between ADA2 and ADA1 structure

The crystal structure of ADA2 (3LGD) and double stranded DNA (1D66) were used for protein-DNA docking using HDOCK server (DOI:10.1093/nar/gkx407). The top scored complex was subjected to steered molecular dynamic simulation using a TIP3P water model in a solvated box under periodic boundary conditions. Minimization was performed using 100 steepest descent steps at 0.02 Å followed by 10 steps of conjugate gradient at 0.02 Å. Equilibration was performed for 100000 steps (time step 1fs) using heater temperature control method. Production was performed for 100000 steps (time step 1fs). Molecular dynamic simulation was performed with UCSF Chimera (*10*). Molecular images were produced using PyMOL Molecular Graphics System, Version 2.0 Schrödinger, LLC.

#### ADA2 WT and ADA2 mutants production and purification

Protein production was performed in 100 ml (small scale screening), and a selection of the mutants in 1 l (large scale). ADA2 plasmid DNA (uniprot Q9NZK5: aa29-511) with a CD33 leader on its N-terminus and a APP- and His6 tag on its C terminus was transfected with the PEI method in HKB11 mammalian cells. Fresh grown cells (1.25 x 10^8^ cells) have been centrifuged, resuspended in 9 ml fresh M11V3 media (Bioconcept, #V3-k) and transferred into 250 ml shake flask. 75 ug DNA and 225 ug PEI have been transferred in 1.75 ml media each and incubated for 5 min in separate. After transfer of PEI to DNA and gentle mixing, both have been incubated for another 15 minutes at RT. Then, DNA- and PEI-Mix has been added to cells and incubated for four hours in shaking incubator (standard cultivation conditions with shaking at 115 rpm at 37°C with 5% CO_2_ and 80% humidity). Subsequently, transfected cells have been fed with 87.5ml M11V3-media, and then incubated for 7 days in shaking incubator. For a 1 l large scale expression with a selection of some mutants, this protocol has been scaled up. Protein was purified from the cell supernatant by ion metal affinity chromatography (IMAC) using the His-tag and rebuffered into PBS, pH7.4. Subsequently, proteins from 1 l large scale production have been polished on preparative SEC (Superdex75 column, GE Healthcare, 28-9893-33). Protein integrity and purity was analyzed by SDS-PAGE and mass spectroscopy (HPLC-ESIMS). The dimerization of the total protein was confirmed by size exclusion chromatography (SEC) and inline multi-angle light scattering (MALS). Human Macrophage colony-stimulating Factor1-Receptor (aa20-512), which comprises nearly the complete extracellular domain of hCSF1-Receptor (aa20-517; UniprotKB-P07333, full length aa1-972) and a His-tag, was expressed and purified as ADA2 and used as a control

#### DNASe and ADA activity of wild type and mutant human recombinant ADA2

ADA2 DNAse of recombinant human ADA2 has been measured by a commercial Fret assay (Thermofisher - AM1970 DNaseAlert). In brief, 6ul ADA2 solution at a final concentration of 10ug/ml was incubated with 3ul DNA-substrate and 40ul of MIB super buffer 2 (Malonic acid: Imidazole: Boric acid in the molar ratios 2:3:3) or CHC super buffer 1 (Citric acid:HEPES:CHES in the molar ratios 2:3:4) (Jena Bioscience CS-332-JBScreen Thermofluor Fundament) at the indicated pH and NaCl concentration. Break down of DNA was measured every 2 mins for 30 minutes with a fluorescent reader (Spectramax, excitation 535 nm and emission 575nm). ADA2 DNAse of small scale purified recombinant human ADA2 mutants has been measured by fret assay at 25μg/ml. the G358R mutant was purified at large scale and its DNAse activity measured at pH 4.8 at different protein concentration and compared to WT human ADA2, human ADA1 and a control protein expressed by the same cell type of hADA2 and purified with the same protocol and machinery. Adenosine Deaminase (ADA) activity has been measured by a commercial assay (Diazyme-DZ117A-K), which is specific for ADA and has no detectable reaction with other nucleosides. 5ul of ADA2 or control proteins were added to 180ul of buffer R1 and incubated 3min at 37°C, then 90ul of buffer R2 was added and the plate was incubated at 37°C for 20min. The enzymatic activity was monitored by the measurement of the OD at 550nm with a temperature controlled fluoresecent reader (Synergy H1 from Biotek).

#### Mass spec analysis of recombinant human ADA2

1 μg of protein was resuspended in 8M urea, 10 mM HEPES (pH 8), 10 mM DTT. Alkylation was performed in the dark for 30 min by adding 55 mM iodoacetamide (IAA). A two-step proteolytic digestion was performed. First, samples were digested at room temperature (RT) with LysC (1:50, w/w) for 3h. Then, they were diluted 1:5 with 50 mM ammoniumbicarbonate (pH 8) and digested with trypsin (1:50, w/w) at RT overnight. The resulting peptide mixtures were acidified and loaded on C18 StageTips. Peptides were eluted with 80% acetonitrile (ACN), dried using a SpeedVac centrifuge (Savant, Concentrator plus, SC 110 A), and resuspended in 2% ACN, 0.1% trifluoroacetic acid (TFA), and 0.5% acetic acid. Peptides were separated on an EASY-nLC 1200 HPLC system (Thermo Fisher Scientific) coupled online to a Q Exactive mass HF spectrometer via a nanoelectrospray source (Thermo Fisher Scientific). Peptides were loaded in buffer A (0.1% formic acid) on in house packed columns (75 μm inner diameter, 50 cm length, and 1.9 μm C18 particles from Dr. Maisch GmbH). Peptides were eluted with a nonlinear 120 min gradient of 5%–60% buffer B (80% ACN, 0.1% formic acid) at a flow rate of 250 nl/min and a column temperature of 50°C. Raw files were analyzed by MaxQuant software (version 1.5.3.54). The mass spectrometry proteomics data have been deposited to the ProteomeXchange Consortium via the PRIDE (*11*) partner repository with the dataset identifier PXD019382.

#### Porcine brain ADA2 purification

A schematic overview of the ADA2 purification steps from 5 kg porcine brains obtained freshly from the local slaughterhouse is shown in Suppl. Fig. S4. The brain was cut into small pieces and homogenized in 0.075 M acetic acid/0.15 M NaCl (1:2 mass/vol.) using a Waring blendor. The homogenate was centrifuged at 10000 g for 10 min. The pH of the supernatant was adjusted by adding 1 M TRIS base until the pH reached 7.6. It was then heat treated at 60°C for 20 min and centrifuged as before. The supernatant was added 20 ml Concanavalin A Sepharose and stirred overnight at 4°C. The slurry was run through a column and washed with PBS. The glycoproteins including ADA2 was eluted using 0.2 M α-methylmannoside in PBS. The protein solution was subsequently loaded onto a 20 ml hydroxyapatite column equilibrated with PBS and the proteins eluted with a stepwise phosphate concentration. The main bulk of ADA2 eluted at 0.05 M phosphate. After dialysis against 0.02 M Tris, pH 7.6 this fraction was applied to a DEAE anion exchange column equilibrated with the same buffer. ADA2 appeared in the run through fraction. It was concentrated and applied to a Sephadex S-200 gel filtration column. The ADA2 isomer with high plasmid DNase activity coeluted with arylsulphatase B with a native mass of about 60 kDa. This fraction was dialyzed against 0.02 M Tris, pH 7.6 and subjected to CM cation exchange chromatography using a continuous NaCl salt gradient. ADA2 eluted at about 0.08 M NaCl and this fraction was dialyzed against 0.02 M Tris, pH 7.6. After dialysis against 0.02 M Tris, pH 7.6 the samples were run in heparin Sepharose chromatography and the ADA2 isomer with high plasmid DNase activity bound strongly to heparin and was eluted at about 0.1 M NaCl. We assessed the purity of the final preparation of ADA2 by SDS/PAGE and MS/MS analyses (Suppl. Fig. 4). The mass spectrometry proteomics data have been deposited to the ProteomeXchange Consortium via the PRIDE (*11*) partner repository with the dataset identifier PXD019373

#### MS/MS-analysis or porcine brain proteins

Gel pieces were subjected to in gel reduction, alkylation, and tryptic digestion using 6 ng/μl trypsin. OMIX C18 tips (Varian, Inc., Palo Alto, CA, USA) was used for sample cleanup and concentration. Peptide mixtures containing 0.1% formic acid were loaded onto a Thermo Fisher Scientific EASY-nLC1000 system and EASY-Spray column (C18, 2μm, 100 Å, 50μm, 15 cm). Peptides were fractionated using a 2-100% acetonitrile gradient in 0.1 % formic acid over 50 min at a flow rate of 250 nl/min. The separated peptides were analysed using a Thermo Scientific Q-Exactive mass spectrometer. Data was collected in data dependent mode using a Top10 method. The Proteome Discoverer 1.4 software was used to generate mgf peak list files. The mgf files was searched against a mammalian database using an in-house Mascot server (Matrix Sciences, UK). Peptide mass tolerances used in the search were 10 ppm, and fragment mass tolerance was 0.02 Da.

#### Site specific analysis of N-glycan structures linked to porcine brain proteins

The N-glycan analysis was carried out essentially as previously described (*12*), but without spectral aligning, due to the purity of the proteins. 1) The peptide sequences of the glycopeptides were determined by calculating the theoretical masses of the tryptic peptides containing NXS/T glycosylation sequons of the target protein. The calculated masses of each of these peptides attached to more than 250 different N-glycan structures were determined and compared to all of the peptide masses obtained from the MS/MS spectra. Those masses that matched were analysed further. 2) The MS/MS-spectra containing glycopeptides were identified by the typical oxonium ions of simple sugars, m/z 163.1 (Hex), m/z 204.1 (HexNAc) and m/z 366.1 (HexHexNAc). Spectra containing mannose-6-phosphorylated glycans were identified by the additional ions, m/z 243.1 (PHex), and its fragmentation ion, m/z 225.1. 3) Next, each individual glycopeptide spectrum was scanned for the presence of fragmentation ions with a similar mass as the deglycosylated peptide, and ions with an additional mass of 203.1, corresponding to linkage with a single HexNAc. 4) The identity of the peptide part was further confirmed by the identification of fragmentation ions corresponding to the calculated y-, b-values. 5) The glycan structure linked to the peptide was deduced from a comparison between its mass as determined from the MS/MS-spectrum and theoretical masses of N-glycan structures. The glycan structures were divided into the following groups, as mentioned in Fig. 2D: SN3, sialylated complex structure with three GlcNAc’s linked to the core; SN2, sialylated complex structure with two GlcNAcs linked to the core; SN1, sialylated complex structure with one GlcNAc linked to the core; NN3, neutral complex structure with three GlcNAcs linked to the core; NN2, neutral complex structure with two GlcNAcs linked to the core; NN1, neutral complex structure with one GlcNAc linked to the core; LMF, core-fucosylated structure with 1-3 mannoses linked to the core chitobiose unit; P2M, bisphosphorylated oligomannosidic structure; P1M, monophosphorylated oligomannosidic structure; HM, oligomannose with 6-9 mannose; MM, oligomannose with 4-5 mannose; LM, oligomannose with 1-3 mannose.

#### Evaluation of porcine brain mediated DNAse activity

The DNA substrate was pcDNA 3.1 with a MEK1 insert (a gift from Ugo Moens, University of Tromso). 10 μl reaction sample containing 25 mM acetic acid buffer of different pH, 0.2 μg plasmid DNA, 0.5 μg BSA and the ADA2 sample, was incubated at room temperature for 10 min, and the reaction stopped by adding 2μl of 1 M Tris base. The DNA fragments were separated by 1.5% agarose gel electrophoresis. RedSafe™ nucleic acid staining solution was used to visualize DNA. For standard assays the pH of the buffer was typically 4.8, corresponding to the pH-optimum for both ADA2 and DNase2.

#### THP1 cells culture, propagation, differentiation and stimulation

THP-1 cells were grown in RPMI 1640 10% FBS, 2 mM L-glutamine. The cells were maintained at a density between 2.5×10^5 cells/ml to 5×10^5 cells/ml, changing the medium every two-three days. Before stimulation the cells were stimulated 6 hrs with 50 nM phorbol 12-myristate 12-acetate [PMA, Sigma P8139], washed twice and stimulated with the following stimuli. THP1 cell lines were pre-activated with PMA [50nM] washed twice with RPMI 10% FCS and subsequently stimulated over night with the following stimuli: Poly-IC (Invivogen https://www.invivogen.com/ cat. num. 31852-29-6, working concentration 1μg/ml, with lipofectamine LTX Invitrogen ratio dsDNA/Lipofectamine, 1μg/μl), LPS (Invivogen https://www.invivogen.com/ cat. num. tlrl-eklps, working concentration 10 ng/ml), Imiquimod (Invivogen https://www.invivogen.com/ cat. num. tlrl-imqs, working concentration 1 μg/ml), ISD70 (VACV 70mer-derived DNA ‘-CCATCAGAAAGAGGTTTAATATTTTTGTGAGACCATCGAAGAGAGAAAGA GATAAAACTTTTTTACGACT-3’, synthetized by SigmaAldrich, working concentration 1μg/ml, complexed with lipofectamine LTX Invitrogen; ratio dsDNA/Lipofectamine, 1μg/1μl).

#### CRISPR-Cas9 knockout of ADA2 and STING in THP1 cells

CRISPR-Cas9 technology was used to generate ADA2 or ADA2/STING knockout THP1 cells. The AATCAAGTTCCCCACGGTGG guide RNA targeting exon 5 of ADA2 and the CTAGCCCCCAAAGGGTCACC targeting exon 4 of STING were get from http://crispr.mit.edu have been cloned into the minimal plasmid pMAX-Crispr with Cas9-2A-eGFP (provided by dr. Bartok E) using Gibson assembly. Briefly the vector was linearized with SwaI, the gRNA was annealed at equimolar ratio with the universal antisense Oligo: GCCTTATTTTAACTTGCTATTTCTAGCTCTAAAAC (annealing cycling: 95°C/5 min, gradient −1°C/30 sec, 80 cycles), the annealed oligos were filled-in with Klenow fragment and finally assembled with the digested vector. Stbl3 competent E. coli were transformed with 2 μl of Assembly. Single colonies were grown in LB (kanamycin), plasmid DNA extracted and Saanger sequenced to confirm the insertion. THP1 cells were electroporated (10 μg/1,25×10^6 cells) using NEON system. The setting of electroporation was the following: 1250V, 1 pulse, 20 ms. The day after electroporation, GFP positive cells have been FACS sorted into 384 well at a density of 0.5 cells/well in RPMI 10% FCS. The growing clones have been screened for gene KO by western blot and/or Sanger sequencing after PCR amplification of the target exon.

#### Lentivirus transduction of THP1-/- cells with ADA2

The ADA2 protein (uniprot Q9NZK5: aa29-511), was clone into pLenti-GFP-puro plasmid. HEK293 cells were transfected with pLenti-ADA2-IRES-GFP-Puro, psPAX2 and pMD2G to produce lentiviral particles. After 48 hrs from transfection the supernatant was transferred to THP1 cells and incubated 16 hr a humidified incubator in an atmosphere of 5-7% CO2. After this time the media containing lentiviral particles was removed and substituted with fresh medium. The efficiency of transduction was monitored measuring GFP expression by flow cytometry (BD Fortessa). Transduced cells were used for experiments starting from three weeks after transduction.

#### SDS-PAGE and Western Blotting

Stimulated or unstimulated cells were transferred into eppendorf tubes and centrifuged at 200 g for 5 min at room temperature. The supernatant was discarded, and the pellet was resuspended in 5 ml cold PBS. After two washing steps of 5 minutes at 4 °C (first: 200 g; second: top speed, 18,000 g), the pellet was resuspended in lysis buffer (200μl per 10 x 106 cells) and 1% PMSF. After 25 minutes incubation on ice, they were sonicated for 20 minutes and finally centrifuged at top speed for 10 minutes at 4° C. The protein concentration was determined with the Pierce™ BCA Protein Assay Kit according to the manufacturer’s protocol. In order to reach a final volume of 30 μL, 5 μL of 6x Laemmli buffer (see below) and the missing amount of ddH20 was added to the appropriate volume of cell lysate, containing 20 −35μg of total protein. The samples were incubated at 70°C for 10 min and afterwards used directly or stored at - 80°C. After the heating step, lysates and protein markers were loaded on a 10% polyacrylamide gel (see below), using a Bio-Rad Mini-PROTEAN Electrophoresis System. The samples were subjected to electrophoresis at 80 V for 30 minutes and subsequently 110 V for another 90 minutes in 1x SDS running buffer. After electrophoresis, the proteins were transferred under wet conditions to a PVDF membrane using the same Bio-Rad Mini-PROTEAN System in 1 x transfer buffer (see below) at 45V for 90 min. Afterwards, the membrane was blocked in blocking buffer (5% powdered milk in Tris-buffered saline (TBS) + 1% Tween20 = TBST) for 1 hour at room temperature. Then, the membrane was incubated with the primary antibodies (anti-CECR1 HPA007888, sigma dilution 1:100, anti-STING, 675902 biolegend dilution 1:500, in blocking buffer plus 0.02% sodium azide in TBST) over night at 4 ° C, followed by 3 washes with TBST. The secondary antibody was also diluted in blocking buffer and the applied for 1 hour at room temperature. After 3 more washes with TBST and another wash with TBS, protein detection was performed using LumiGLO^®^ chemiluminescent substrate according to the manufacturer’s instructions. After an HRP-inactivation step with 10% sodium azide in blocking buffer (1 hour at room temperature), anti-ß-Actin was stained subsequently as a loading control, following the same protocol.

#### Detection of ADA2, pIRF3, and secreted IFN in THP-1 cells after stimulation

THP-1 cells were stimulated with ISD, cGAMP, poly-IC, Imiquimod, ODN-2006 and lipopolysaccharide (LPS) as a control. ADA2, pIRF3, and βactin was detected in cell lysates by automated western blot system (WES, ProteinSimple) using rabbit anti ADA2/CECR1 (Novusbio; # NBP1-89238), rabbit anti pIRF3 (phospho S386; Abcam # ab76493) and anti β-actin (Sigma; # A5441).

#### cGAMP extraction and LC-MS/MS detection

One million cells have been lysed with 400uL cold methanol (−80°C) (Sigma-Aldrich, Germany) containing [15N5]-2,3-cGAMP (Novartis, Switzerland) as internal standard (IS). The collected extract was dried down and resuspended in 100uL 80% acetonitrile (Sigma-Aldrich, Germany). 10uL of this solution was injected for LC-MS/MS analysis. The QTRAP 6500 mass spectrometer (SCIEX, Framingham, MA) was coupled to liquid chromatography (Thermo Fisher Scientific, Waltham, MA). 2,3-cGAMP and IS were detected in the positive ion mode with the mass transition 675.1/524.0 for cGAMP and 680.1/528.9, respectively. All results are given as normalized peak area ratio of cGAMP against IS.

#### HEK-Blue IFN-α/β report cells culture and propagation and IFN-α/β secretion measurement

HEK-Blue IFN-α/β report system was used to detect Type-I IFN, following the protocol described by the manufacturer. Briefly, cells were maintained in DMEM containing Blasticidin (30μg/ml) and Zeocin (100 μg/ml). For detection of Type-I interferon, 180 μl of cell suspension containing 50000 cells were transferred into each well of a 96 flat well plate. The cells were incubated in the incubator for 2 hr. Subsequently 20 μl of supernatant were transferred into each well and incubated 24 hr. IFN-alfa was used as positive control. After incubation 20 μl of cell culture medium were mixed with 180 μl of QUANTI-Blue and type-I IFN detected by spectrophotometer at 660 nm.

**Fig. S1.**
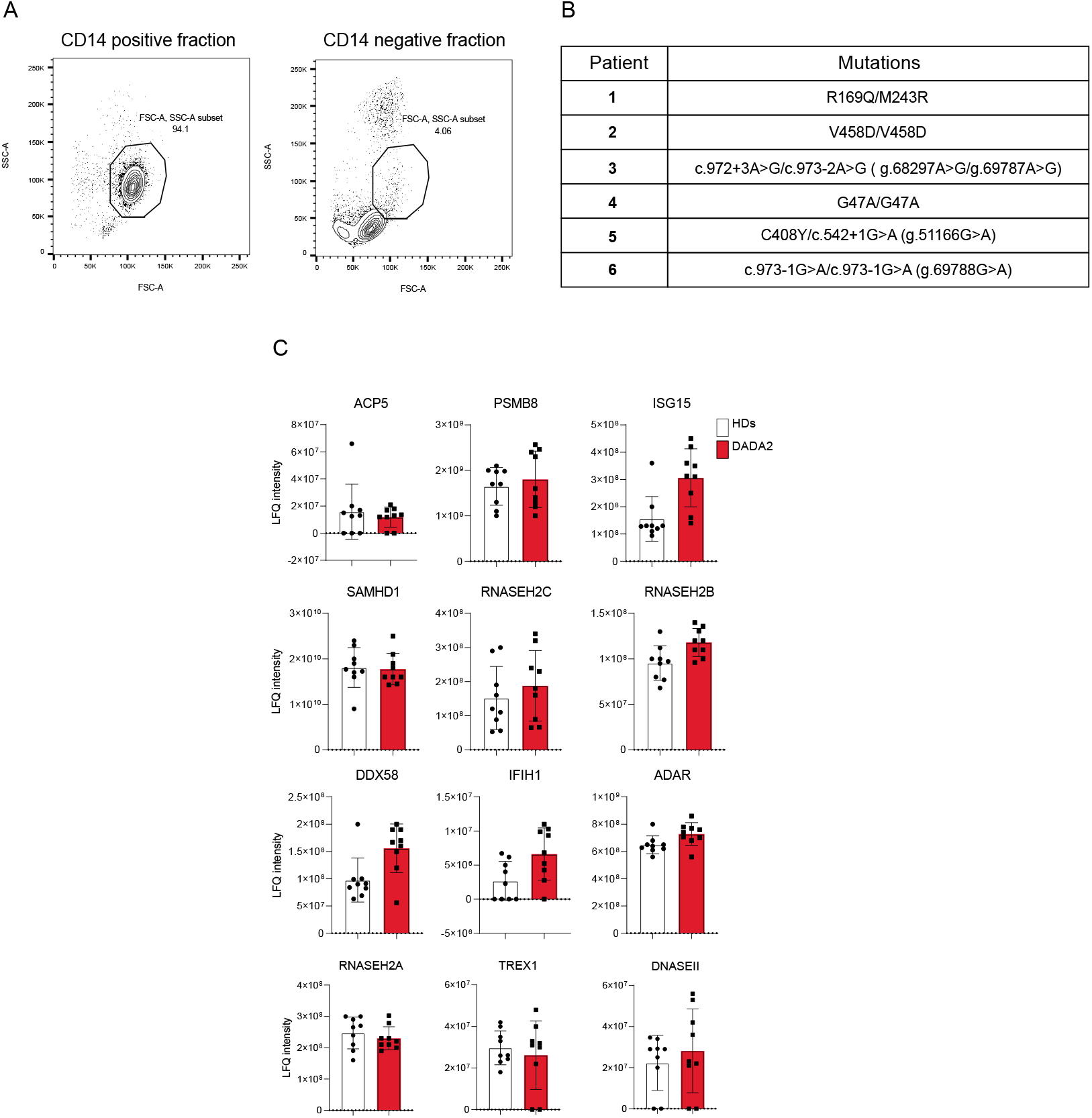
Proteomes of ADA2 deficient monocytes cells. (A) FACS analysis of monocytes purification after magnetic activated cell sorting. (B) *ADA2* mutations in DADA2 patients used for monocytes isolation and monocyte proteomes analysis. (C) LFQ intensity of selected proteins known to be mutated in patients with Type I Interferonopathies (*13*).

**Fig. S2.**
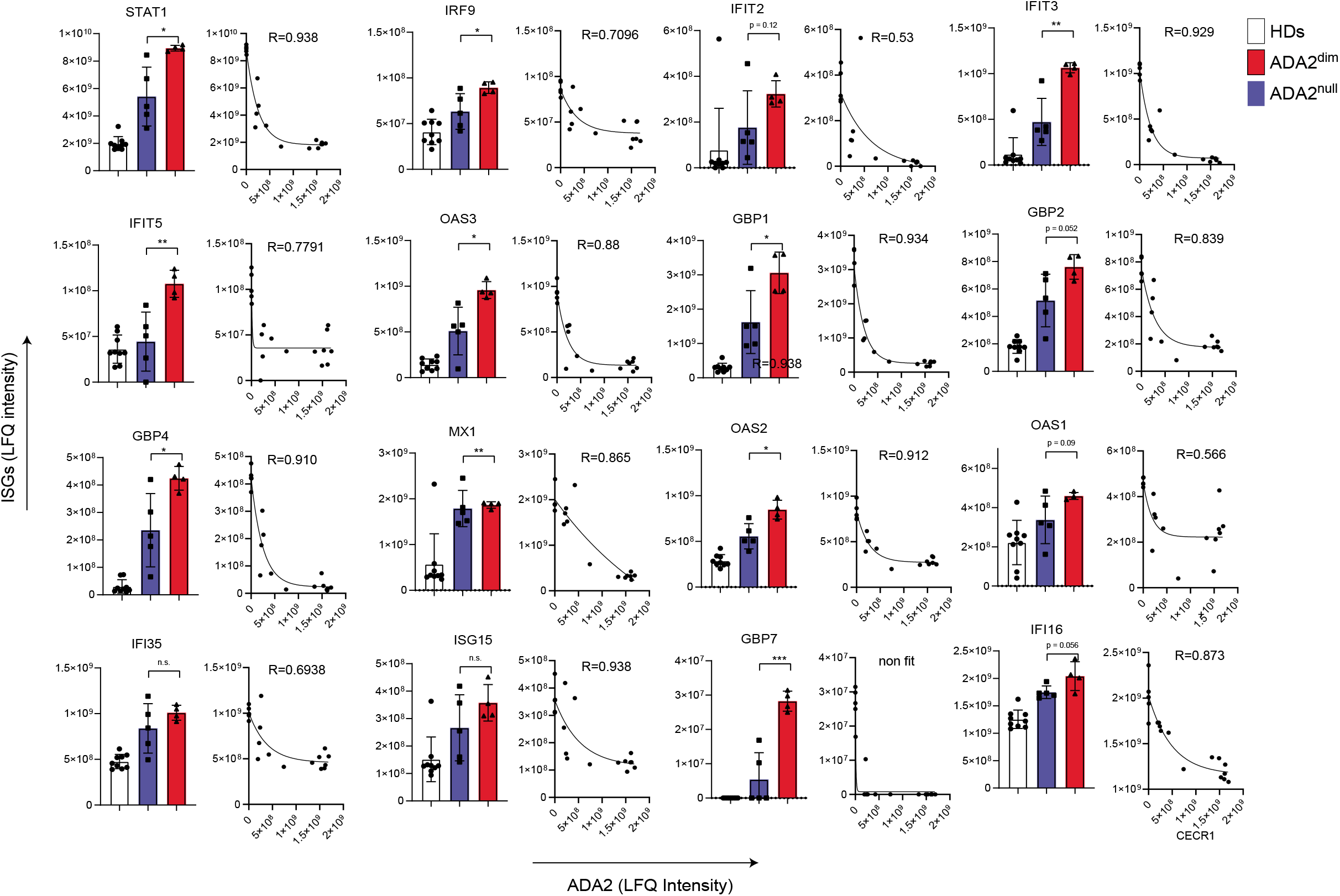
Correlation between ADA2 and ISGs protein levels in proteomes of DADA2 and healthy controls. Left panels: LFQ intensity of different interferon induced genes was compared between healthy controls, DADA2 patients with undetectable ADA2 (ADA2^null^) and DADA2 patients with residual ADA2 protein (ADA2^dim^) (line over the bars represent the statistical comparison, unpaired t.test was used for comparison, , * = p <= 0.01, ** = p <= 0.001, *** = p <= 0.0001. Right panels: correlation between ADA2 and ISG protein levels. The lines show the result of the nonlinear regression and the texts the obtained R squared values.

**Fig. S3.**
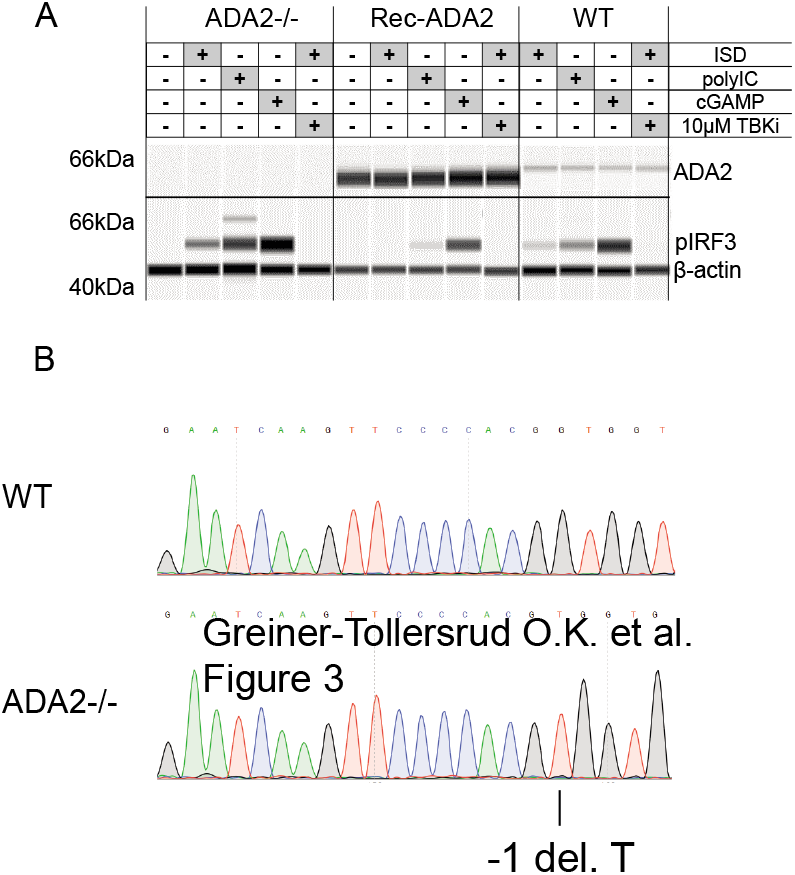
Generation of ADA2^-/-^ monocyte THP1 cell line. (A) ADA2 Western Blot analysis of ADA2, pIRF3, and beta-actin expression in WT *ADA2^-/-^* and *ADA2^-/-^* lentivirus transduced with recombinant human ADA2 THP1 cells. (B) Sanger sequencing of *ADA2* gDNA region (exon 5) targeted by the guide CTAGCCCCCAAAGGGTCACC in the selected exon 5 of ADA2 of the *ADA2*^-/-^ THP1 selected cell clone.

**Fig. S4.**
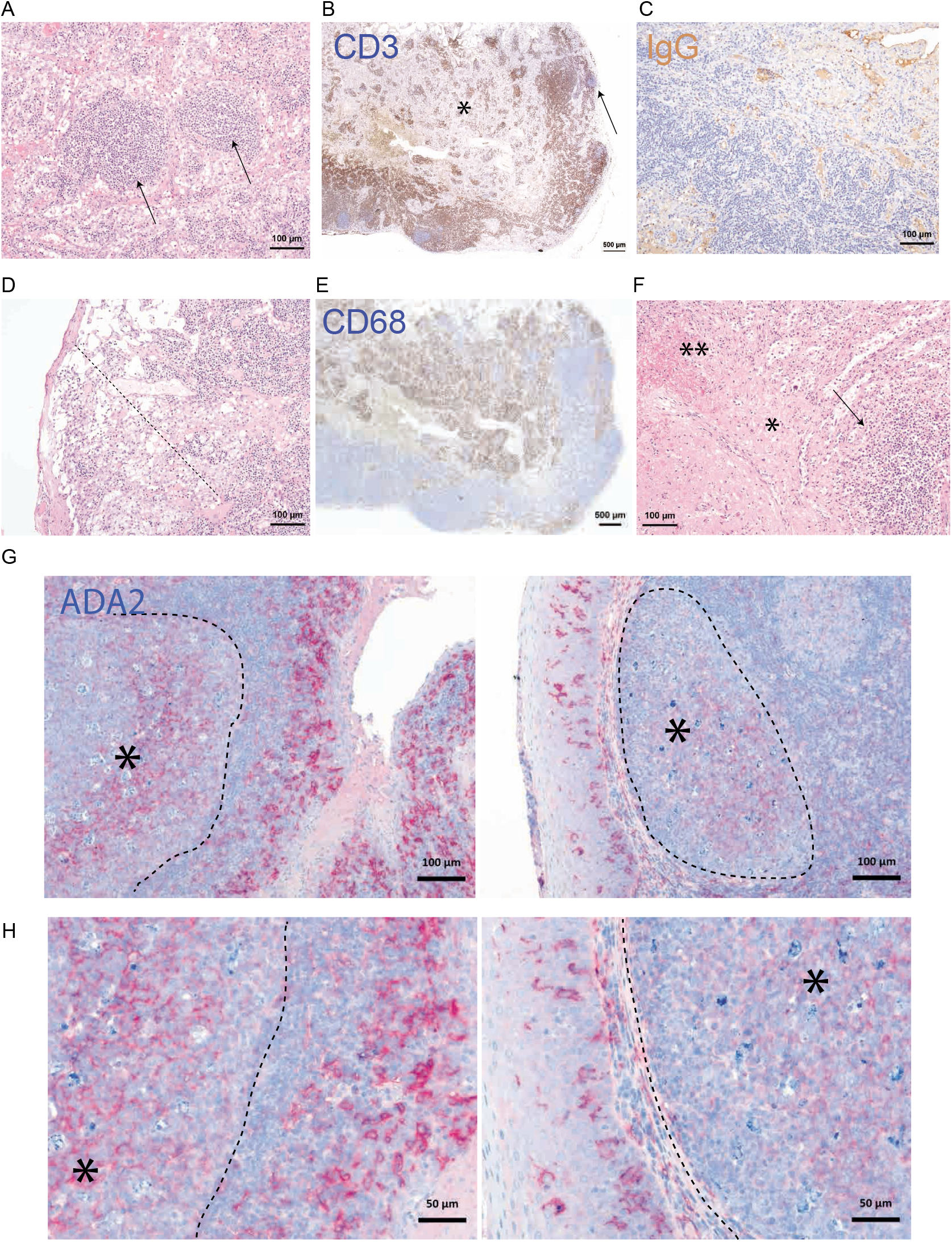
Histopathology of DADA2 secondary lymphoid organs and expression of ADA2 in tingible body macrophages. (A-E) Lymph node; (A) Few follicles (pointed by arrowhead) withou germinal centers. (B) CD3 staining displays normally distributed T-cells (arrowhead points to B-cell follicle with lower density of CD3+ T-cells). (C) Lack of IgG+ plasma cells. Serum is IgG positive due to replacement therapy. The expanded stroma of the lymph node is highlighted by asterisk. (D) Severely ectatic subcapsular sinus (dotted line) with multiple macrophages, most of them foam cells histiocytosis. (E) CD68 displays the multiple macrophages of the sinus F) Spleen: Intact periarteriolar lymphocytic sheath (pointed by arrowhead) next to an infarction (kollagenous scar tissue marked by asterisk, fibrin marked by double asterisk). (G-H) Immunohistochemistry of human tonsil showing CD14 (red) and ADA2 (blue); dotted line show germinal centers with cytoplasmic ADA2 positivity (asterisk, exemplary) in tingible body macrophages; CD14 (red) highlights the follicular dendritic cell meshwork (10x original magnification). In higher magnifications (bottom panels), CD14^+^ dendritic cells in the crypt epithelium (left panel) and surface epithelium (right panel) show few small ADA2 positive intracytoplasmic granules (highlighted by arrowheads, exemplary; 20x original magnification).

**Fig. S5.**
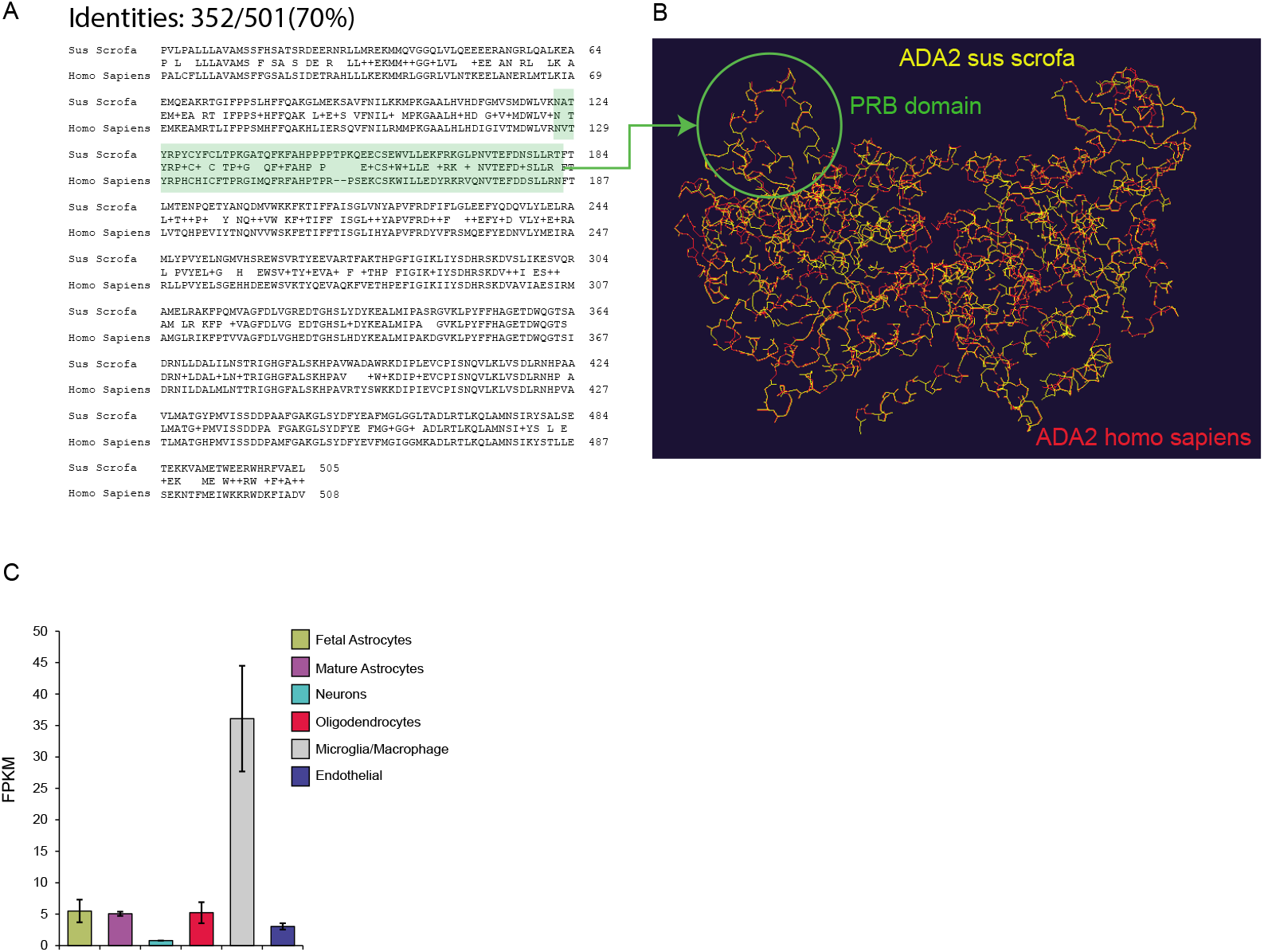
Porcine brain ADA2. (A) Alignment between Sus Scrofa and Homo Sapiens protein ADA2 sequences showing percentage of identity. (B) Comparison between Sus Scrofa (yellow) and Homo Sapiens (Red) ADA2 structure. Highlighted in gray is the PRB domain. Homology between human and porcine ADA2. RNA expression of ADA2 in different cell subtypes isolated from human brain, values are expressed in fragments per kilobase of exon (FPKM) (*14*).

**Fig. S6.**
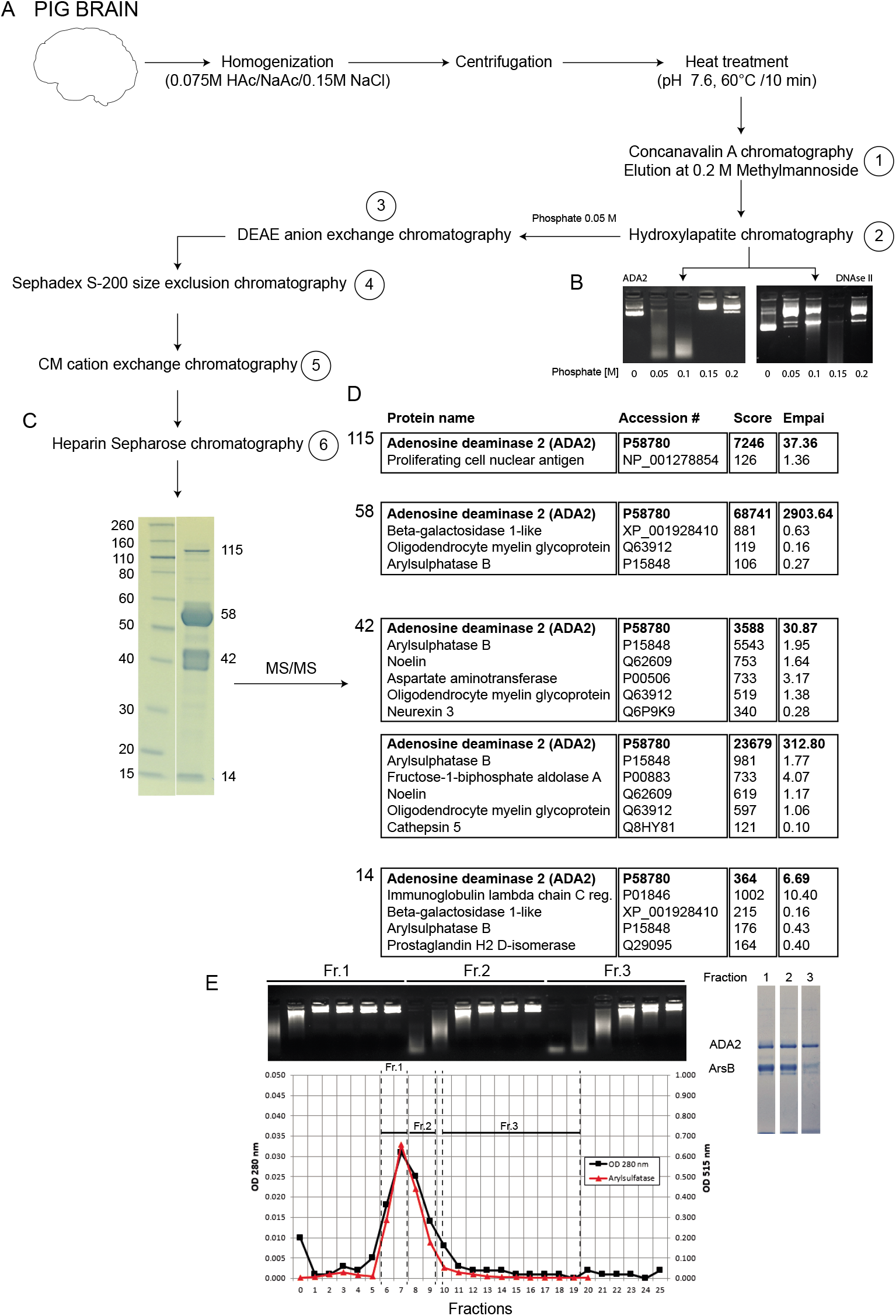
Purification of endogenous ADA2. The figure shows the experimental workflow used for the purification of ADA2 from porcine brain, as described in the methods section. (A) Glycoproteins of a heat-treated porcine brain homogenate were enriched by concanavalin A affinity chromatography (step 1). ADA2 was eluted at 0.05 M phosphate on hydroxyapatite chromatography (step 2), which was instrumental to remove any traces of DNase2. As shown in (B), DNase2 dependent DNase activity eluted on hydroxyapatite chromatography at 0.15 M phospate, while ADA2-mediated DNase activity at 0.05 M. The picture in (B) shows degradation of plasmid DNA after incubation with various fractions from hydroxylapatite chromatography; on left side left side porcine brain proteins after step 1, and on right side a commercial preparation of partially purified DNAse2 from porcine spleen. The DNase containing fraction from step 2 was dialyzed against 0.02 M Tris, pH 7.6 and loaded onto a DEAE column (step 3). The DNase activity appeared in the run through, was concentrated and loaded onto a gel filtration column (step 4). The DNase activity coeluted with arylsuphatase B (ArsB) with a native mass of 60 kDa. After dialysis against 0.02 M Tris, pH 7.6, the eluate was loaded onto a CM Sepharose cation exchange column (step 5). The DNase activity eluted at about 0.08 M NaCl, and SDS/PAGE revealed only two major proteins, ADA2 and ArsB. After dialysis against 0.02 M Tris, pH 7.6, the partially purified DNase was subjected to heparin Sepharose chromatography (step 6). The DNase activity eluted at high salt concentration. (D) MS/MS analyses of tryptic peptides from each band detected on an overloaded SDS/PAGE after one round of heparin Sepharose chromatography (E) revealed that each of the detectable bands contained predominantly ADA2-sequences, except a band of 42 kDa, where the highest score was caused by heavy chain of ArsB (D). Repeated rounds on heparin Sepharose removed all traces of ArsB, which bound with less affinity than the highly DNase active ADA2-isoform (F).

**Fig. S7.**
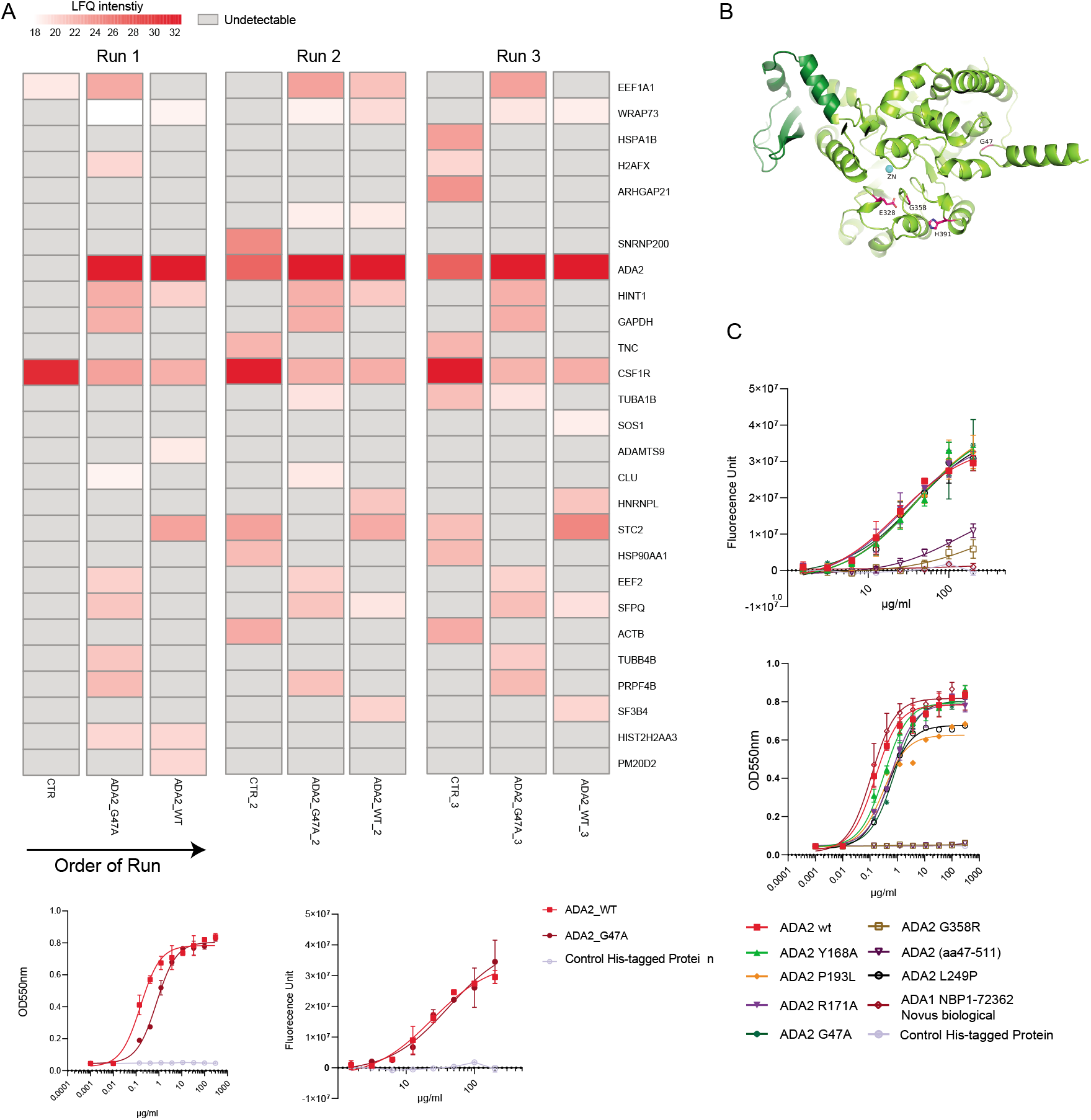
(A) Mass spectrometry analysis made in triplicates of the WT recombinant human ADA2, G47A ADA2 mutant and His/tag purification control and (lower panels) DNAse and ADA activity of the analyzed proteins. (B) The crystal structure of human ADA2 (PDB ID code 3LGD) was used for modelling mutations. Modelling and structural analysis were performed using the program Coot (*15*). Structural illustrations were prepared with PyMOL (www.pymol.org). Comparison between the ADA1 (PDB entries: 3IAR) (34), and ADA2 (3LGD) structure showing the absence of a ~40 amino acids insertion (aa 111-147), corresponding to the PRB domain in ADA1 in comparison to ADA2. (C) Evaluation of DNase and ADA activities of 8 recombinant ADA2 mutants, ADA1 and Control His tag protein purified with IMAC and SEC (see Suppl. Table.

**Table S1.**
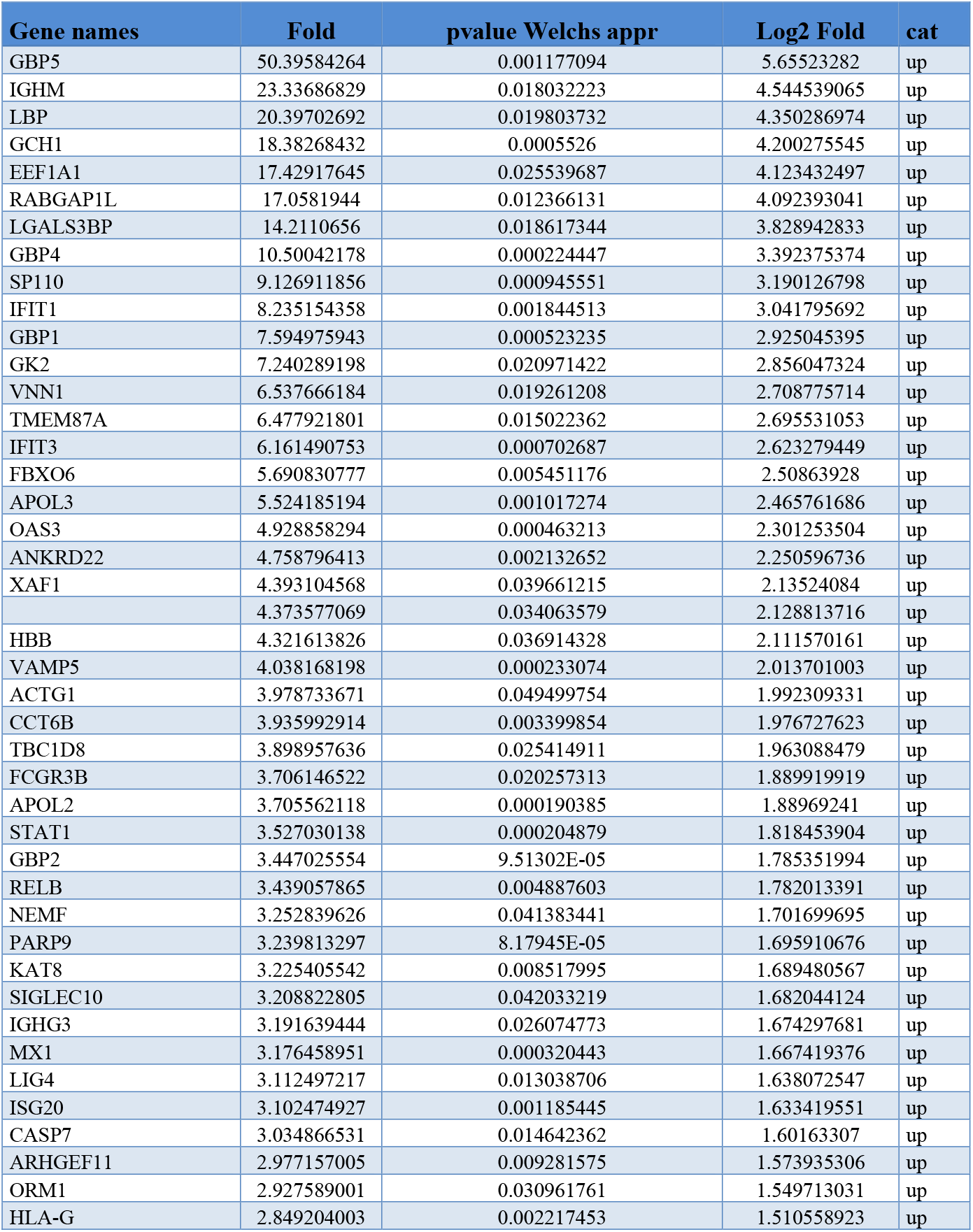
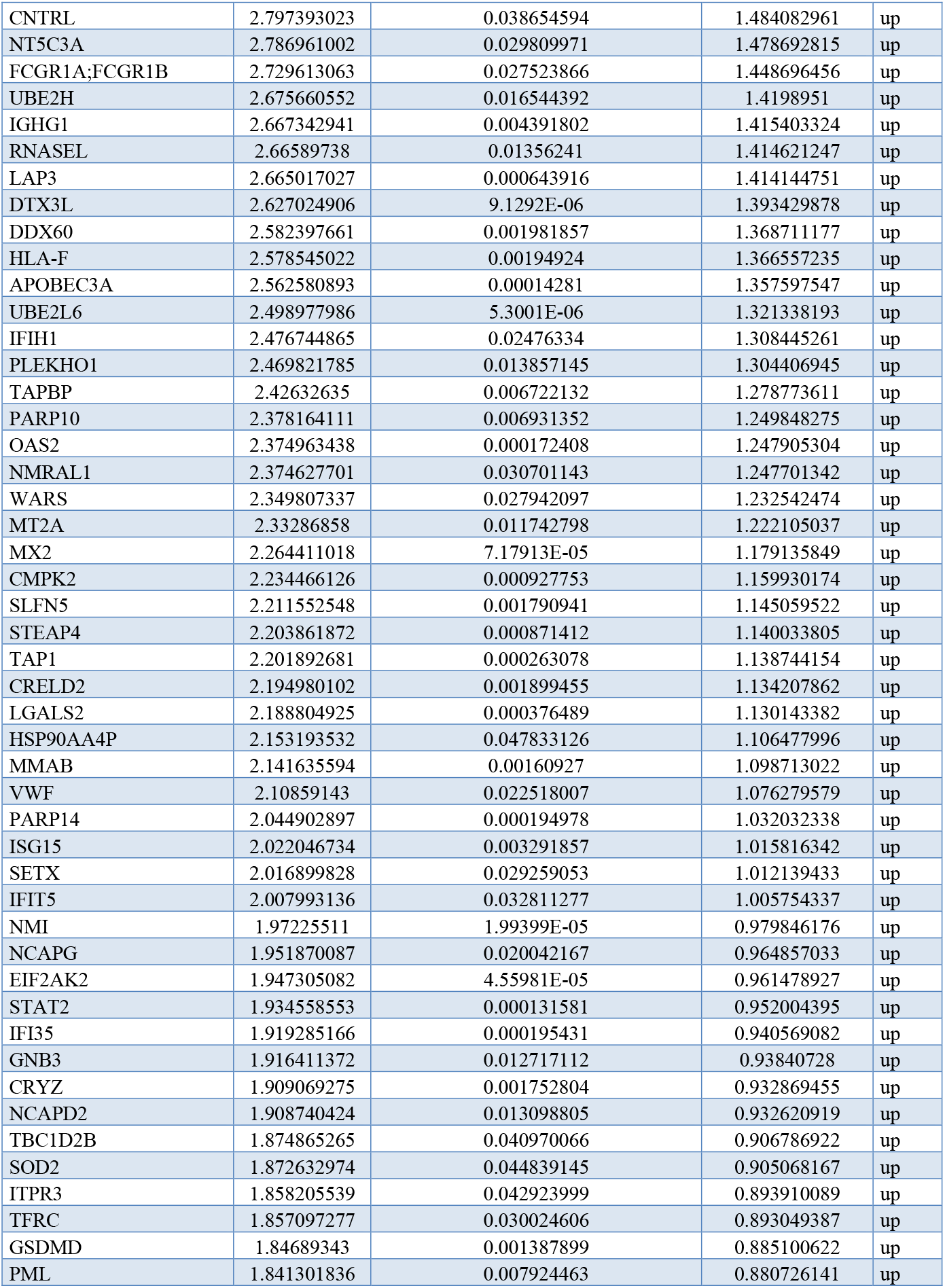
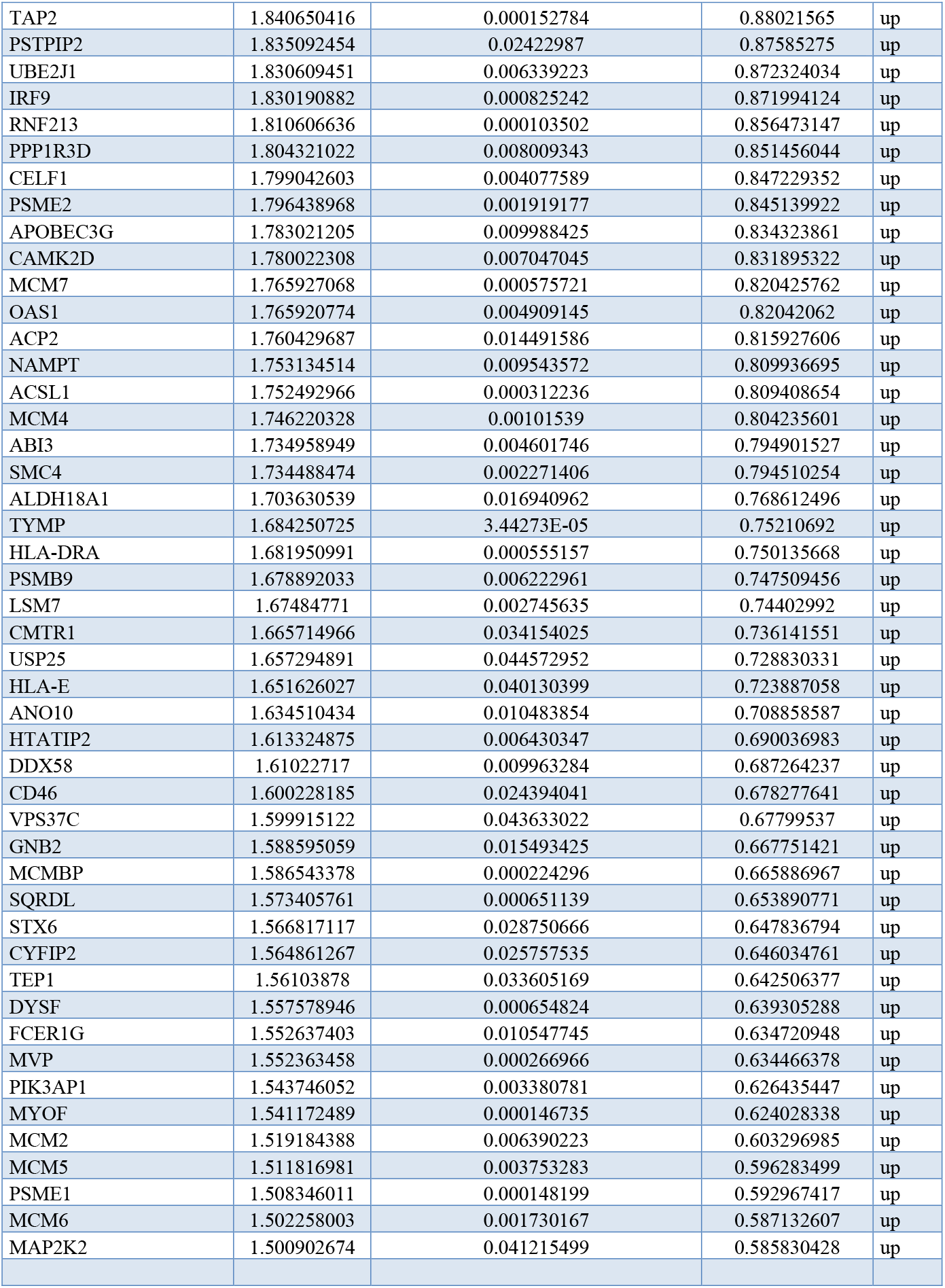
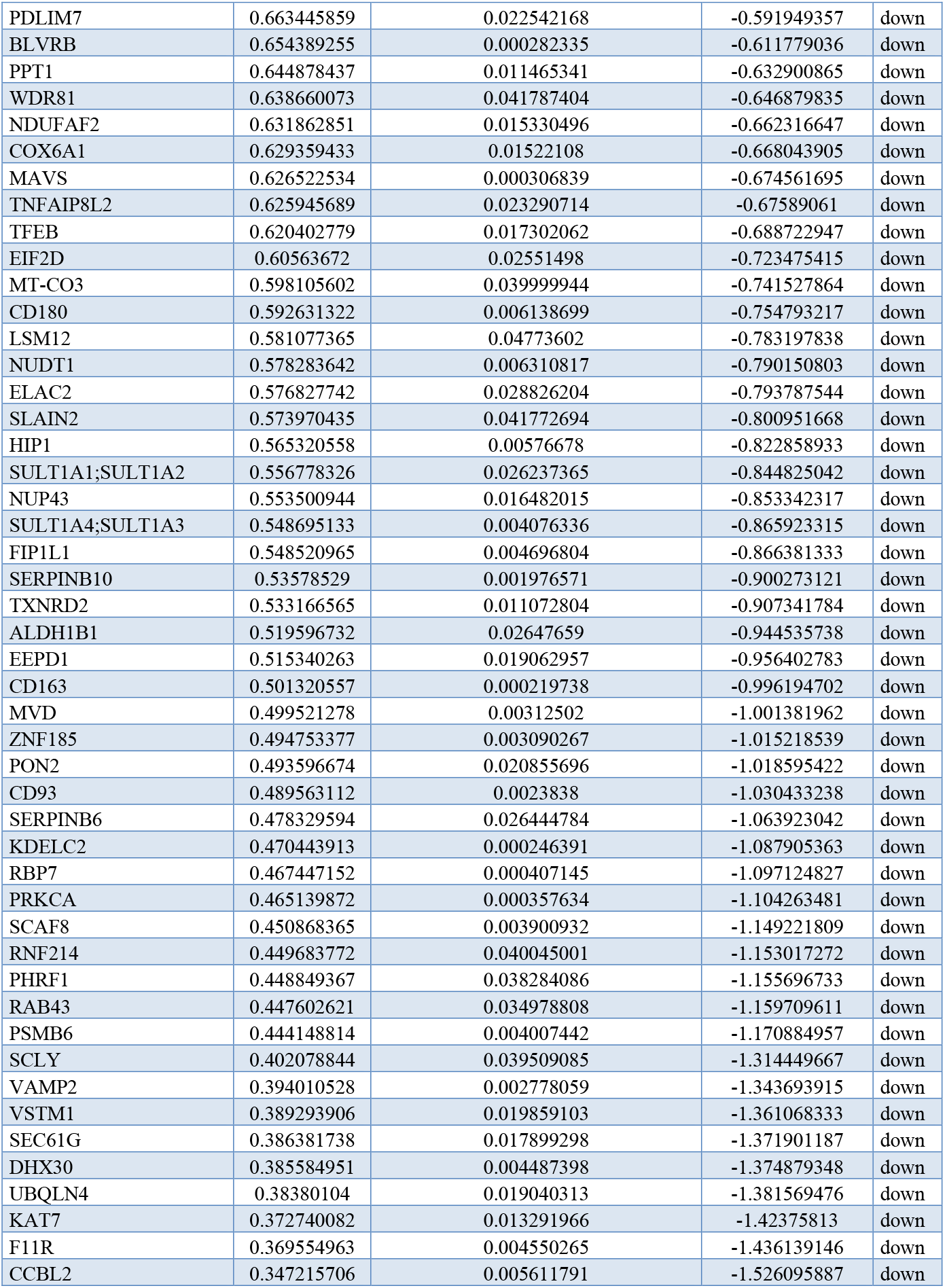
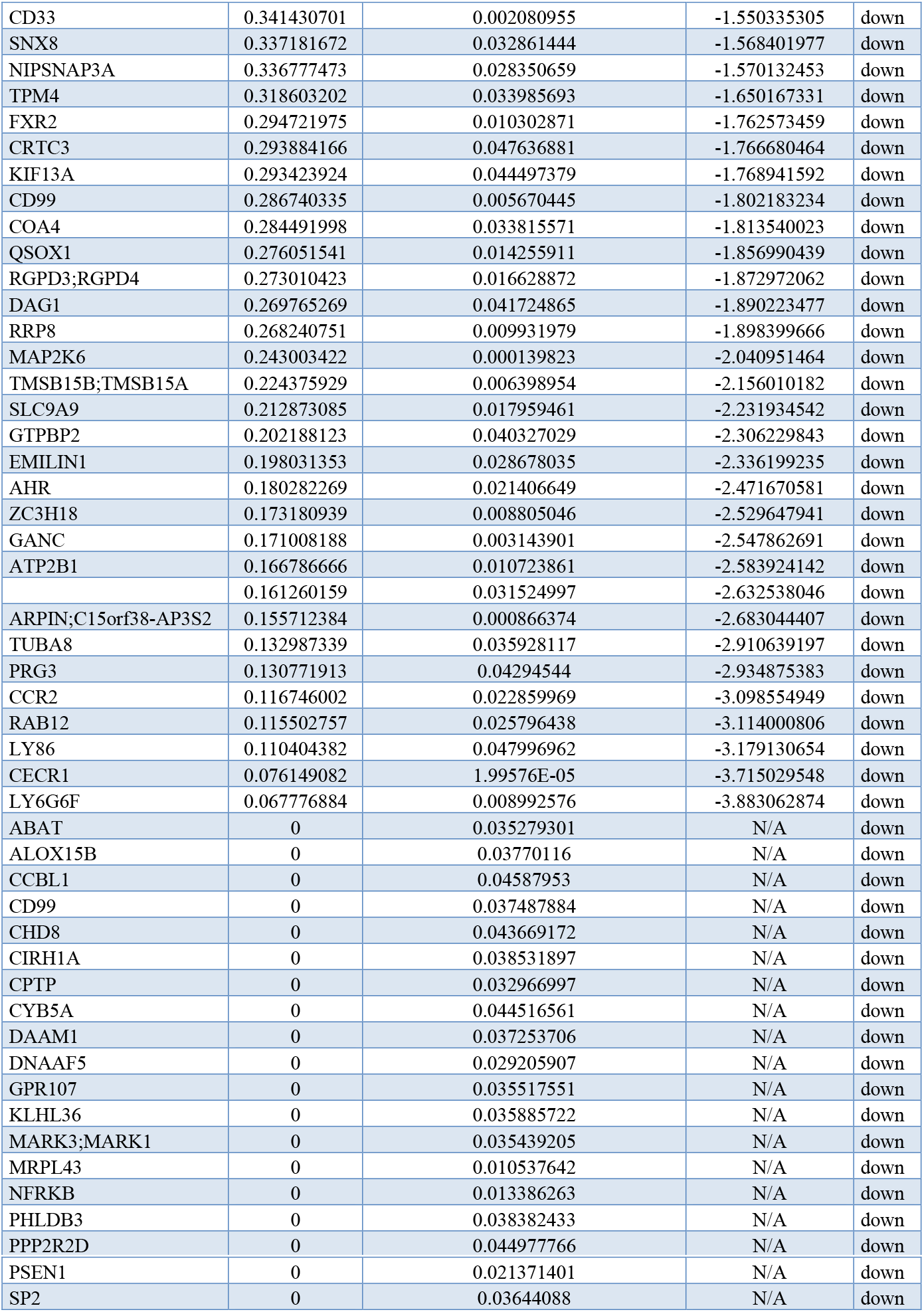
Proteins significantly up or downregulated in proteomes from DADA2 monocytes in comparison with healthy donors controls

| Gene names | Fold | pvalue Welchs appr | Log2 Fold | cat |
| --- | --- | --- | --- | --- |
| GBP5 | 50.39584264 | 0.001177094 | 5.65523282 | up |
| IGHM | 23.33686829 | 0.018032223 | 4.544539065 | up |
| LBP | 20.39702692 | 0.019803732 | 4.350286974 | up |
| GCH1 | 18.38268432 | 0.0005526 | 4.200275545 | up |
| EEF1A1 | 17.42917645 | 0.025539687 | 4.123432497 | up |
| RABGAP1L | 17.0581944 | 0.012366131 | 4.092393041 | up |
| LGALS3BP | 14.2110656 | 0.018617344 | 3.828942833 | up |
| GBP4 | 10.50042178 | 0.000224447 | 3.392375374 | up |
| SP110 | 9.126911856 | 0.000945551 | 3.190126798 | up |
| IFIT1 | 8.235154358 | 0.001844513 | 3.041795692 | up |
| GBP1 | 7.594975943 | 0.000523235 | 2.925045395 | up |
| GK2 | 7.240289198 | 0.020971422 | 2.856047324 | up |
| VNN1 | 6.537666184 | 0.019261208 | 2.708775714 | up |
| TMEM87A | 6.477921801 | 0.015022362 | 2.695531053 | up |
| IFIT3 | 6.161490753 | 0.000702687 | 2.623279449 | up |
| FBXO6 | 5.690830777 | 0.005451176 | 2.50863928 | up |
| APOL3 | 5.524185194 | 0.001017274 | 2.465761686 | up |
| OAS3 | 4.928858294 | 0.000463213 | 2.301253504 | up |
| ANKRD22 | 4.758796413 | 0.002132652 | 2.250596736 | up |
| XAF1 | 4.393104568 | 0.039661215 | 2.13524084 | up |
|  | 4.373577069 | 0.034063579 | 2.128813716 | up |
| HBB | 4.321613826 | 0.036914328 | 2.111570161 | up |
| VAMP5 | 4.038168198 | 0.000233074 | 2.013701003 | up |
| ACTG1 | 3.978733671 | 0.049499754 | 1.992309331 | up |
| CCT6B | 3.935992914 | 0.003399854 | 1.976727623 | up |
| TBC1D8 | 3.898957636 | 0.025414911 | 1.963088479 | up |
| FCGR3B | 3.706146522 | 0.020257313 | 1.889919919 | up |
| APOL2 | 3.705562118 | 0.000190385 | 1.88969241 | up |
| STAT1 | 3.527030138 | 0.000204879 | 1.818453904 | up |
| GBP2 | 3.447025554 | 9.51302E-05 | 1.785351994 | up |
| RELB | 3.439057865 | 0.004887603 | 1.782013391 | up |
| NEMF | 3.252839626 | 0.041383441 | 1.701699695 | up |
| PARP9 | 3.239813297 | 8.17945E-05 | 1.695910676 | up |
| KAT8 | 3.225405542 | 0.008517995 | 1.689480567 | up |
| SIGLEC10 | 3.208822805 | 0.042033219 | 1.682044124 | up |
| IGHG3 | 3.191639444 | 0.026074773 | 1.674297681 | up |
| MX1 | 3.176458951 | 0.000320443 | 1.667419376 | up |
| LIG4 | 3.112497217 | 0.013038706 | 1.638072547 | up |
| ISG20 | 3.102474927 | 0.001185445 | 1.633419551 | up |
| CASP7 | 3.034866531 | 0.014642362 | 1.60163307 | up |
| ARHGEF11 | 2.977157005 | 0.009281575 | 1.573935306 | up |
| ORM1 | 2.927589001 | 0.030961761 | 1.549713031 | up |
| HLA-G | 2.849204003 | 0.002217453 | 1.510558923 | up |
| CNTRL | 2.797393023 | 0.038654594 | 1.484082961 | up |
| NT5C3A | 2.786961002 | 0.029809971 | 1.478692815 | up |
| FCGR1A;FCGR1B | 2.729613063 | 0.027523866 | 1.448696456 | up |
| UBE2H | 2.675660552 | 0.016544392 | 1.4198951 | up |
| IGHG1 | 2.667342941 | 0.004391802 | 1.415403324 | up |
| RNASEL | 2.66589738 | 0.01356241 | 1.414621247 | up |
| LAP3 | 2.665017027 | 0.000643916 | 1.414144751 | up |
| DTX3L | 2.627024906 | 9.1292E-06 | 1.393429878 | up |
| DDX60 | 2.582397661 | 0.001981857 | 1.368711177 | up |
| HLA-F | 2.578545022 | 0.00194924 | 1.366557235 | up |
| APOBEC3A | 2.562580893 | 0.00014281 | 1.357597547 | up |
| UBE2L6 | 2.498977986 | 5.3001E-06 | 1.321338193 | up |
| IFIH1 | 2.476744865 | 0.02476334 | 1.308445261 | up |
| PLEKHO1 | 2.469821785 | 0.013857145 | 1.304406945 | up |
| TAPBP | 2.42632635 | 0.006722132 | 1.278773611 | up |
| PARP10 | 2.378164111 | 0.006931352 | 1.249848275 | up |
| OAS2 | 2.374963438 | 0.000172408 | 1.247905304 | up |
| NMRAL1 | 2.374627701 | 0.030701143 | 1.247701342 | up |
| WARS | 2.349807337 | 0.027942097 | 1.232542474 | up |
| MT2A | 2.33286858 | 0.011742798 | 1.222105037 | up |
| MX2 | 2.264411018 | 7.17913E-05 | 1.179135849 | up |
| CMPK2 | 2.234466126 | 0.000927753 | 1.159930174 | up |
| SLFN5 | 2.211552548 | 0.001790941 | 1.145059522 | up |
| STEAP4 | 2.203861872 | 0.000871412 | 1.140033805 | up |
| TAP1 | 2.201892681 | 0.000263078 | 1.138744154 | up |
| CRELD2 | 2.194980102 | 0.001899455 | 1.134207862 | up |
| LGALS2 | 2.188804925 | 0.000376489 | 1.130143382 | up |
| HSP90AA4P | 2.153193532 | 0.047833126 | 1.106477996 | up |
| MMAB | 2.141635594 | 0.00160927 | 1.098713022 | up |
| VWF | 2.10859143 | 0.022518007 | 1.076279579 | up |
| PARP14 | 2.044902897 | 0.000194978 | 1.032032338 | up |
| ISG15 | 2.022046734 | 0.003291857 | 1.015816342 | up |
| SETX | 2.016899828 | 0.029259053 | 1.012139433 | up |
| IFIT5 | 2.007993136 | 0.032811277 | 1.005754337 | up |
| NMI | 1.97225511 | 1.99399E-05 | 0.979846176 | up |
| NCAPG | 1.951870087 | 0.020042167 | 0.964857033 | up |
| EIF2AK2 | 1.947305082 | 4.55981E-05 | 0.961478927 | up |
| STAT2 | 1.934558553 | 0.000131581 | 0.952004395 | up |
| IFI35 | 1.919285166 | 0.000195431 | 0.940569082 | up |
| GNB3 | 1.916411372 | 0.012717112 | 0.93840728 | up |
| CRYZ | 1.909069275 | 0.001752804 | 0.932869455 | up |
| NCAPD2 | 1.908740424 | 0.013098805 | 0.932620919 | up |
| TBC1D2B | 1.874865265 | 0.040970066 | 0.906786922 | up |
| SOD2 | 1.872632974 | 0.044839145 | 0.905068167 | up |
| ITPR3 | 1.858205539 | 0.042923999 | 0.893910089 | up |
| TFRC | 1.857097277 | 0.030024606 | 0.893049387 | up |
| GSDMD | 1.84689343 | 0.001387899 | 0.885100622 | up |
| PML | 1.841301836 | 0.007924463 | 0.880726141 | up |
| TAP2 | 1.840650416 | 0.000152784 | 0.88021565 | up |
| PSTPIP2 | 1.835092454 | 0.02422987 | 0.87585275 | up |
| UBE2J1 | 1.830609451 | 0.006339223 | 0.872324034 | up |
| IRF9 | 1.830190882 | 0.000825242 | 0.871994124 | up |
| RNF213 | 1.810606636 | 0.000103502 | 0.856473147 | up |
| PPP1R3D | 1.804321022 | 0.008009343 | 0.851456044 | up |
| CELF1 | 1.799042603 | 0.004077589 | 0.847229352 | up |
| PSME2 | 1.796438968 | 0.001919177 | 0.845139922 | up |
| APOBEC3G | 1.783021205 | 0.009988425 | 0.834323861 | up |
| CAMK2D | 1.780022308 | 0.007047045 | 0.831895322 | up |
| MCM7 | 1.765927068 | 0.000575721 | 0.820425762 | up |
| OAS1 | 1.765920774 | 0.004909145 | 0.82042062 | up |
| ACP2 | 1.760429687 | 0.014491586 | 0.815927606 | up |
| NAMPT | 1.753134514 | 0.009543572 | 0.809936695 | up |
| ACSL1 | 1.752492966 | 0.000312236 | 0.809408654 | up |
| MCM4 | 1.746220328 | 0.00101539 | 0.804235601 | up |
| ABI3 | 1.734958949 | 0.004601746 | 0.794901527 | up |
| SMC4 | 1.734488474 | 0.002271406 | 0.794510254 | up |
| ALDH18A1 | 1.703630539 | 0.016940962 | 0.768612496 | up |
| TYMP | 1.684250725 | 3.44273E-05 | 0.75210692 | up |
| HLA-DRA | 1.681950991 | 0.000555157 | 0.750135668 | up |
| PSMB9 | 1.678892033 | 0.006222961 | 0.747509456 | up |
| LSM7 | 1.67484771 | 0.002745635 | 0.74402992 | up |
| CMTR1 | 1.665714966 | 0.034154025 | 0.736141551 | up |
| USP25 | 1.657294891 | 0.044572952 | 0.728830331 | up |
| HLA-E | 1.651626027 | 0.040130399 | 0.723887058 | up |
| ANO10 | 1.634510434 | 0.010483854 | 0.708858587 | up |
| HTATIP2 | 1.613324875 | 0.006430347 | 0.690036983 | up |
| DDX58 | 1.61022717 | 0.009963284 | 0.687264237 | up |
| CD46 | 1.600228185 | 0.024394041 | 0.678277641 | up |
| VPS37C | 1.599915122 | 0.043633022 | 0.67799537 | up |
| GNB2 | 1.588595059 | 0.015493425 | 0.667751421 | up |
| MCMBP | 1.586543378 | 0.000224296 | 0.665886967 | up |
| SQRDL | 1.573405761 | 0.000651139 | 0.653890771 | up |
| STX6 | 1.566817117 | 0.028750666 | 0.647836794 | up |
| CYFIP2 | 1.564861267 | 0.025757535 | 0.646034761 | up |
| TEP1 | 1.56103878 | 0.033605169 | 0.642506377 | up |
| DYSF | 1.557578946 | 0.000654824 | 0.639305288 | up |
| FCER1G | 1.552637403 | 0.010547745 | 0.634720948 | up |
| MVP | 1.552363458 | 0.000266966 | 0.634466378 | up |
| PIK3AP1 | 1.543746052 | 0.003380781 | 0.626435447 | up |
| MYOF | 1.541172489 | 0.000146735 | 0.624028338 | up |
| MCM2 | 1.519184388 | 0.006390223 | 0.603296985 | up |
| MCM5 | 1.511816981 | 0.003753283 | 0.596283499 | up |
| PSME1 | 1.508346011 | 0.000148199 | 0.592967417 | up |
| MCM6 | 1.502258003 | 0.001730167 | 0.587132607 | up |
| MAP2K2 | 1.500902674 | 0.041215499 | 0.585830428 | up |
| PDLIM7 | 0.663445859 | 0.022542168 | -0.591949357 | down |
| BLVRB | 0.654389255 | 0.000282335 | -0.611779036 | down |
| PPT1 | 0.644878437 | 0.011465341 | -0.632900865 | down |
| WDR81 | 0.638660073 | 0.041787404 | -0.646879835 | down |
| NDUFAF2 | 0.631862851 | 0.015330496 | -0.662316647 | down |
| COX6A1 | 0.629359433 | 0.01522108 | -0.668043905 | down |
| MAVS | 0.626522534 | 0.000306839 | -0.674561695 | down |
| TNFAIP8L2 | 0.625945689 | 0.023290714 | -0.67589061 | down |
| TFEB | 0.620402779 | 0.017302062 | -0.688722947 | down |
| EIF2D | 0.60563672 | 0.02551498 | -0.723475415 | down |
| MT-CO3 | 0.598105602 | 0.039999944 | -0.741527864 | down |
| CD180 | 0.592631322 | 0.006138699 | -0.754793217 | down |
| LSM12 | 0.581077365 | 0.04773602 | -0.783197838 | down |
| NUDT1 | 0.578283642 | 0.006310817 | -0.790150803 | down |
| ELAC2 | 0.576827742 | 0.028826204 | -0.793787544 | down |
| SLAIN2 | 0.573970435 | 0.041772694 | -0.800951668 | down |
| HIP1 | 0.565320558 | 0.00576678 | -0.822858933 | down |
| SULT1A1;SULT1A2 | 0.556778326 | 0.026237365 | -0.844825042 | down |
| NUP43 | 0.553500944 | 0.016482015 | -0.853342317 | down |
| SULT1A4;SULT1A3 | 0.548695133 | 0.004076336 | -0.865923315 | down |
| FIP1L1 | 0.548520965 | 0.004696804 | -0.866381333 | down |
| SERPINB10 | 0.53578529 | 0.001976571 | -0.900273121 | down |
| TXNRD2 | 0.533166565 | 0.011072804 | -0.907341784 | down |
| ALDH1B1 | 0.519596732 | 0.02647659 | -0.944535738 | down |
| EEPD1 | 0.515340263 | 0.019062957 | -0.956402783 | down |
| CD163 | 0.501320557 | 0.000219738 | -0.996194702 | down |
| MVD | 0.499521278 | 0.00312502 | -1.001381962 | down |
| ZNF185 | 0.494753377 | 0.003090267 | -1.015218539 | down |
| PON2 | 0.493596674 | 0.020855696 | -1.018595422 | down |
| CD93 | 0.489563112 | 0.0023838 | -1.030433238 | down |
| SERPINB6 | 0.478329594 | 0.026444784 | -1.063923042 | down |
| KDELC2 | 0.470443913 | 0.000246391 | -1.087905363 | down |
| RBP7 | 0.467447152 | 0.000407145 | -1.097124827 | down |
| PRKCA | 0.465139872 | 0.000357634 | -1.104263481 | down |
| SCAF8 | 0.450868365 | 0.003900932 | -1.149221809 | down |
| RNF214 | 0.449683772 | 0.040045001 | -1.153017272 | down |
| PHRF1 | 0.448849367 | 0.038284086 | -1.155696733 | down |
| RAB43 | 0.447602621 | 0.034978808 | -1.159709611 | down |
| PSMB6 | 0.444148814 | 0.004007442 | -1.170884957 | down |
| SCLY | 0.402078844 | 0.039509085 | -1.314449667 | down |
| VAMP2 | 0.394010528 | 0.002778059 | -1.343693915 | down |
| VSTM1 | 0.389293906 | 0.019859103 | -1.361068333 | down |
| SEC61G | 0.386381738 | 0.017899298 | -1.371901187 | down |
| DHX30 | 0.385584951 | 0.004487398 | -1.374879348 | down |
| UBQLN4 | 0.38380104 | 0.019040313 | -1.381569476 | down |
| KAT7 | 0.372740082 | 0.013291966 | -1.42375813 | down |
| F11R | 0.369554963 | 0.004550265 | -1.436139146 | down |
| CCBL2 | 0.347215706 | 0.005611791 | -1.526095887 | down |
| CD33 | 0.341430701 | 0.002080955 | -1.550335305 | down |
| SNX8 | 0.337181672 | 0.032861444 | -1.568401977 | down |
| NIPSNAP3A | 0.336777473 | 0.028350659 | -1.570132453 | down |
| TPM4 | 0.318603202 | 0.033985693 | -1.650167331 | down |
| FXR2 | 0.294721975 | 0.010302871 | -1.762573459 | down |
| CRTC3 | 0.293884166 | 0.047636881 | -1.766680464 | down |
| KIF13A | 0.293423924 | 0.044497379 | -1.768941592 | down |
| CD99 | 0.286740335 | 0.005670445 | -1.802183234 | down |
| COA4 | 0.284491998 | 0.033815571 | -1.813540023 | down |
| QSOX1 | 0.276051541 | 0.014255911 | -1.856990439 | down |
| RGPD3;RGPD4 | 0.273010423 | 0.016628872 | -1.872972062 | down |
| DAG1 | 0.269765269 | 0.041724865 | -1.890223477 | down |
| RRP8 | 0.268240751 | 0.009931979 | -1.898399666 | down |
| MAP2K6 | 0.243003422 | 0.000139823 | -2.040951464 | down |
| TMSB15B;TMSB15A | 0.224375929 | 0.006398954 | -2.156010182 | down |
| SLC9A9 | 0.212873085 | 0.017959461 | -2.231934542 | down |
| GTPBP2 | 0.202188123 | 0.040327029 | -2.306229843 | down |
| EMILIN1 | 0.198031353 | 0.028678035 | -2.336199235 | down |
| AHR | 0.180282269 | 0.021406649 | -2.471670581 | down |
| ZC3H18 | 0.173180939 | 0.008805046 | -2.529647941 | down |
| GANC | 0.171008188 | 0.003143901 | -2.547862691 | down |
| ATP2B1 | 0.166786666 | 0.010723861 | -2.583924142 | down |
|  | 0.161260159 | 0.031524997 | -2.632538046 | down |
| ARPIN;C15orf38-AP3S2 | 0.155712384 | 0.000866374 | -2.683044407 | down |
| TUBA8 | 0.132987339 | 0.035928117 | -2.910639197 | down |
| PRG3 | 0.130771913 | 0.04294544 | -2.934875383 | down |
| CCR2 | 0.116746002 | 0.022859969 | -3.098554949 | down |
| RAB12 | 0.115502757 | 0.025796438 | -3.114000806 | down |
| LY86 | 0.110404382 | 0.047996962 | -3.179130654 | down |
| CECR1 | 0.076149082 | 1.99576E-05 | -3.715029548 | down |
| LY6G6F | 0.067776884 | 0.008992576 | -3.883062874 | down |
| ABAT | 0 | 0.035279301 | N/A | down |
| ALOX15B | 0 | 0.03770116 | N/A | down |
| CCBL1 | 0 | 0.04587953 | N/A | down |
| CD99 | 0 | 0.037487884 | N/A | down |
| CHD8 | 0 | 0.043669172 | N/A | down |
| CIRH1A | 0 | 0.038531897 | N/A | down |
| CPTP | 0 | 0.032966997 | N/A | down |
| CYB5A | 0 | 0.044516561 | N/A | down |
| DAAM1 | 0 | 0.037253706 | N/A | down |
| DNAAF5 | 0 | 0.029205907 | N/A | down |
| GPR107 | 0 | 0.035517551 | N/A | down |
| KLHL36 | 0 | 0.035885722 | N/A | down |
| MARK3;MARK1 | 0 | 0.035439205 | N/A | down |
| MRPL43 | 0 | 0.010537642 | N/A | down |
| NFRKB | 0 | 0.013386263 | N/A | down |
| PHLDB3 | 0 | 0.038382433 | N/A | down |
| PPP2R2D | 0 | 0.044977766 | N/A | down |
| PSEN1 | 0 | 0.021371401 | N/A | down |
| SP2 | 0 | 0.03644088 | N/A | down |

**Table S.2:** Comparison of the main clinical signs and symptoms observed respectively in ADA1 deficiency, ADA2 deficiency and DNAseII deficiency.

| Symptom | ADA1 deficiency | ADA2 deficiency | DNaseII deficiency |
| --- | --- | --- | --- |
| Age of onset | <6 months | Variable | Variable |
| Absence of lymph node germinal centers | Frequent | N.R. | N.R. |
| Absent tonsils | Frequent | N.R. | N.R. |
| Allergy | Frequent | Variable | Variable |
| B– lymphocytopenia | Frequent | Frequent | Variable |
| Decrease in t cell count | Frequent | Rare | Rare |
| Chronic Diarrhea | Frequent | Variable | Variable |
| Failure to thrive | Frequent | Variable | Variable |
| Increased circulating total IgE level | Frequent | Variable | Variable |
| Diffuse rash | Frequent | Rare | Rare |
| Livedo racemosa | Rare | Frequent | Frequent |
| Purpura | Rare | Frequent | Variable |
| Recurrent otitis media | Frequent | Variable | Variable |
| Recurrent pneumonia | Frequent | Variable | Variable |
| Recurrent respiratory tract infections | Frequent | Variable | Variable |
| Sinusitis | Frequent | Variable | Variable |
| Vasculitis | Rare | Frequent | Frequent |
| Pancytopenia | Rare | Frequent | Frequent |
| Anemia | Rare | Frequent | Frequent |
| Digital ischemia | Rare | Frequent | Frequent |
| Ischemic stroke | Rare | Frequent | Frequent |
| Glomerulopathy | Rare | Frequent | Frequent |
| Arthropathy | Rare | Rare | Frequent |

**Table S.3:** Expression yields of ADA2 mutants. For screening in small scale expression each mutant construct was produced in 100ml final volume. Final expression yield after IMAC purification are noted in mg. For the large-scale expression, several structural mutants were selected but only mutants with yield above 0.4 mg in small scale expression. Each of these mutants was produced in 1L scale, purified via IMAC and SEC. Final yield in mg.* G47A, was not purified by SEC as it already had a high monomer content after IMAC purification.

| Construct | Small scale | Large scale#<br>(IMAC only/IMAC+SEC) |
| --- | --- | --- |
| M243R | 0.03 |  |
| G321E | 0.04 |  |
| Domain aa29-465 | 0.05 |  |
| Domain aa29-191 | 0.08 |  |
| Domain aa96-511 | 0.09 |  |
| L311R | 0.10 |  |
| Deletion aa170-177 | 0.11 |  |
| T129P | 0.12 |  |
| V458D | 0.13 |  |
| aa132-511 | 0.13 |  |
| I93T | 0.14 |  |
| F178S | 0.18 |  |
| M445K | 0.31 |  |
| L249P | 0.31 | 5.4 / 2.9 |
| G47R | 0.45 |  |
| G47A |  | 12 / * |
| P193L | 0.47 | 5.2 / 2.7 |
| E328K | 0.6 |  |
| G358R | 0.92 | 7.7 / 1.5 |
| Y168A | 0.96 | 13.2 / 10.6 |
| Domain aa47-511 | 0.97 | 12.1 / 4.6 |
| P344L | 1.23 |  |
| K170A | 2.1 |  |
| ADA2 Wt | 2.61 | 28.6 / 7.6 |
| R171A | 2.8 | 33.4 / 29.1 |

